# Integrating multiple genomic technologies to investigate an outbreak of carbapenemase-producing *Enterobacter hormaechei*

**DOI:** 10.1101/172536

**Authors:** Leah W. Roberts, Patrick N. A. Harris, Brian M. Forde, Nouri L. Ben Zakour, Mitchell Stanton-Cook, Elizabeth Catchpoole, Minh-Duy Phan, Hanna E. Sidjabat, Haakon Bergh, Claire Heney, Jayde A. Gawthorne, Jeffrey Lipman, Anthony Allworth, Kok-Gan Chan, Teik Min Chong, Wai-Fong Yin, Mark A. Schembri, David L. Paterson, Scott A. Beatson

## Abstract

Carbapenem-resistant Enterobacteriaceae (CRE) represent one of the most urgent threats to human health posed by antibiotic resistant bacteria. *Enterobacter hormaechei* and other members of the *Enterobacter cloacae* complex are the most commonly encountered *Enterobacter* spp. within clinical settings, responsible for numerous outbreaks and ultimately poorer patient outcomes. Here we applied three complementary whole genome sequencing (WGS) technologies to characterise a hospital cluster of *bla*_IMP-4_ carbapenemase-producing *E. hormaechei*.

In response to a suspected CRE outbreak in 2015 within an Intensive Care Unit (ICU)/Burns Unit in a Brisbane tertiary referral hospital we used Illumina sequencing to determine that all outbreak isolates were sequence type (ST)90 and near-identical at the core genome level. Comparison to publicly available data unequivocally linked all 10 isolates to a 2013 isolate from the same ward, confirming the hospital environment as the most likely original source of infection in the 2015 cases. No clonal relationship was found to IMP-4-producing isolates identified from other local hospitals. However, using Pacific Biosciences long-read sequencing we were able to resolve the complete context of the *bla*_IMP-4_ gene, which was found to be on a large IncHI2 plasmid carried by all IMP-4-producing isolates. Continued surveillance of the hospital environment was carried out using Oxford Nanopore long-read sequencing, which was able to rapidly resolve the true relationship of subsequent isolates to the initial outbreak. Shotgun metagenomic sequencing of environmental samples also found evidence of ST90 *E. hormaechei* and the IncHI2 plasmid within the hospital plumbing.

Overall, our strategic application of three WGS technologies provided an in-depth analysis of the outbreak, including the transmission dynamics of a carbapenemase-producing *E. hormaechei* cluster, identification of possible hospital reservoirs and the full context of *bla*_IMP-4_ on a multidrug resistant IncHI2 plasmid that appears to be widely distributed in Australia.

## Introduction

Carbapenem antibiotics have become the mainstay of therapy for serious infections caused by multidrug resistant (MDR) Gram-negative bacteria, especially for strains expressing extended-spectrum β-lactamase (ESBL) or AmpC-type enzymes^1^. Increased use has driven resistance to carbapenems and the emergence of carbapenemase-producing Enterobacteriaceae (CPE) and carbapenem-resistant Enterobacteriaceae (CRE), which include common enteric species such as *Escherichia coli, Klebsiella pneumoniae* and *Enterobacter* spp.^2^.

Before 2005, an estimated 99.9% of Enterobacteriaceae were susceptible to carbapenems^3^. However, the isolation of CRE has since increased dramatically and these organisms are now reported in all WHO health regions^4^. The mortality rates for CRE infections are reported to be as high as 48%^5^, and resistance to last-line antibiotics used in lieu of carbapenems, such as colistin, has also emerged^6^.

Resistance to carbapenems in Enterobacteriaceae occurs via a range of mechanisms. Of greatest concern is the acquisition of genes encoding carbapenemases^7^. This most frequently occurs via transfer of mobile genetic elements (MGE), such as plasmids, occasionally carrying multiple β-lactamases co-located with other resistance determinants, rendering these strains MDR or extensively drug-resistant (XDR)^8^. Australia has experienced low rates of CRE ^9^, although sporadic introduction of *K. pneumoniae* carbapenemase (KPC)^10^ and New Delhi metallo-β-lactamase (NDM) 11 has been reported, including significant nosocomial outbreaks^12^. The most frequently encountered carbapenemase in Australia is *bla*_IMP-4_, particularly in *Enterobacter* spp.^13^. IMP-producing *Enterobacter* spp. have caused occasional outbreaks within intensive care or burns units in Australian hospitals^14–16^.

Here, we describe the use of whole genome sequencing (WGS) to investigate an outbreak of IMP-4-producing *Enterobacter hormaechei* within an Intensive Care Unit (ICU) and Burns facility.

### Clinical case report

Two patients in mid 2015 were transferred from regional Queensland hospitals to the ICU with burn injuries sustained from the same accident (Figure 1). *E. cloacae* complex was cultured from the endotracheal tube (ETT) of patients 1 and 2 on day 6 and 8 of admission, respectively. Both *E. cloacae* complex isolates were confirmed as MDR by phenotypic testing used in the diagnostic setting (Table 1). Real-time PCR amplification of *bla*_IMP-4_ confirmed their status as carbapenemase-producers. Both of these patients were previously well, with no prior hospital admission or contact with healthcare facilities. Neither had been resident or hospitalized overseas for more than 20 years.

**Figure 1:**
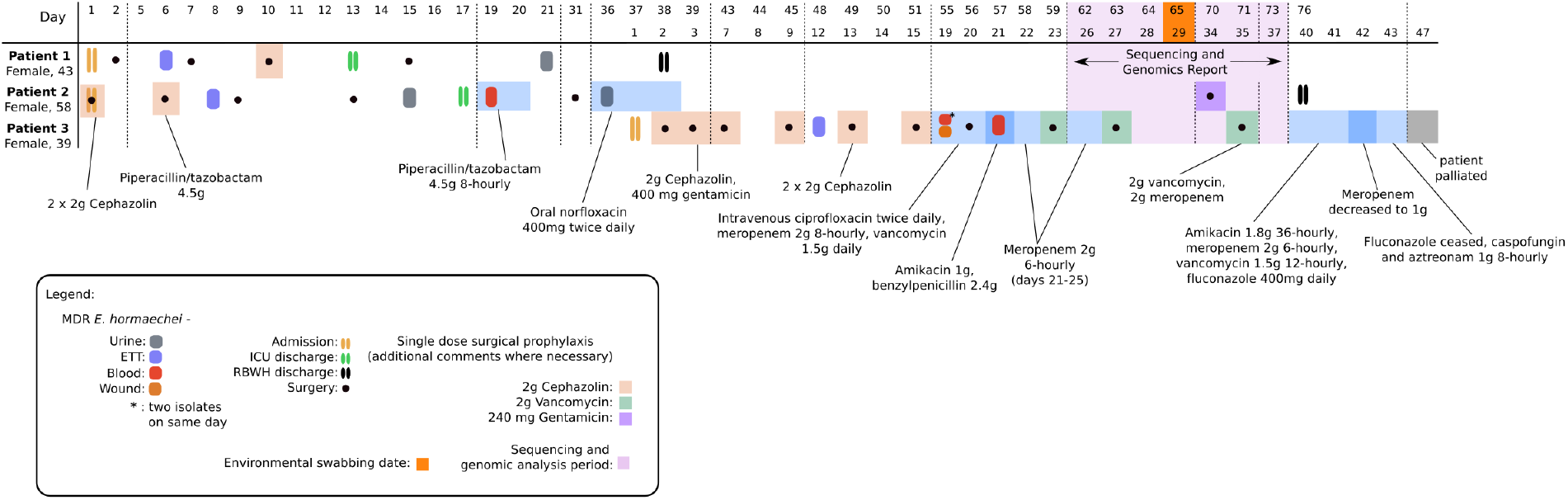
RBWH clinical case study outline. Three burns patients were admitted to the RBWH ICU ward in mid 2015. Patient 1 (Female, 43-years-old) and Patient 2 (Female, 58-years-old) were admitted on the same day. Subsequent to admission, both patients developed carbapenem-resistant *E. hormaechei* infections, with two samples taken from patient 1 (source = ETT [purple] and urine [grey]), and 4 samples taken from patient 2 (source = ETT [purple], urine [grey], and blood [red]). Patient 3 (Female, 39-years-old) was admitted 37 days after the patient 1 and 2 had been admitted and after they had been discharged from the ICU. Patient 3 also developed infection due to a carbapenem-resistant *E. cloacae* infection, and had 4 samples taken from ETT (purple), blood (red) and wound sites (orange). After intensive antibiotic and antifungal treatment, the patient was palliated on day 47 of ICU admission. Sequencing and genomics analysis of all 10 isolates was undertaken following confirmation of all three patients being infected with *bla*_IMP-4_-producing *E. hormaechei* (period shown in purple shading). Environmental swabbing was undertaken 65 days after the initial admission of patient 1 and 2, and 29 days after the admission of patient 3 (orange square).

**Table 1:**
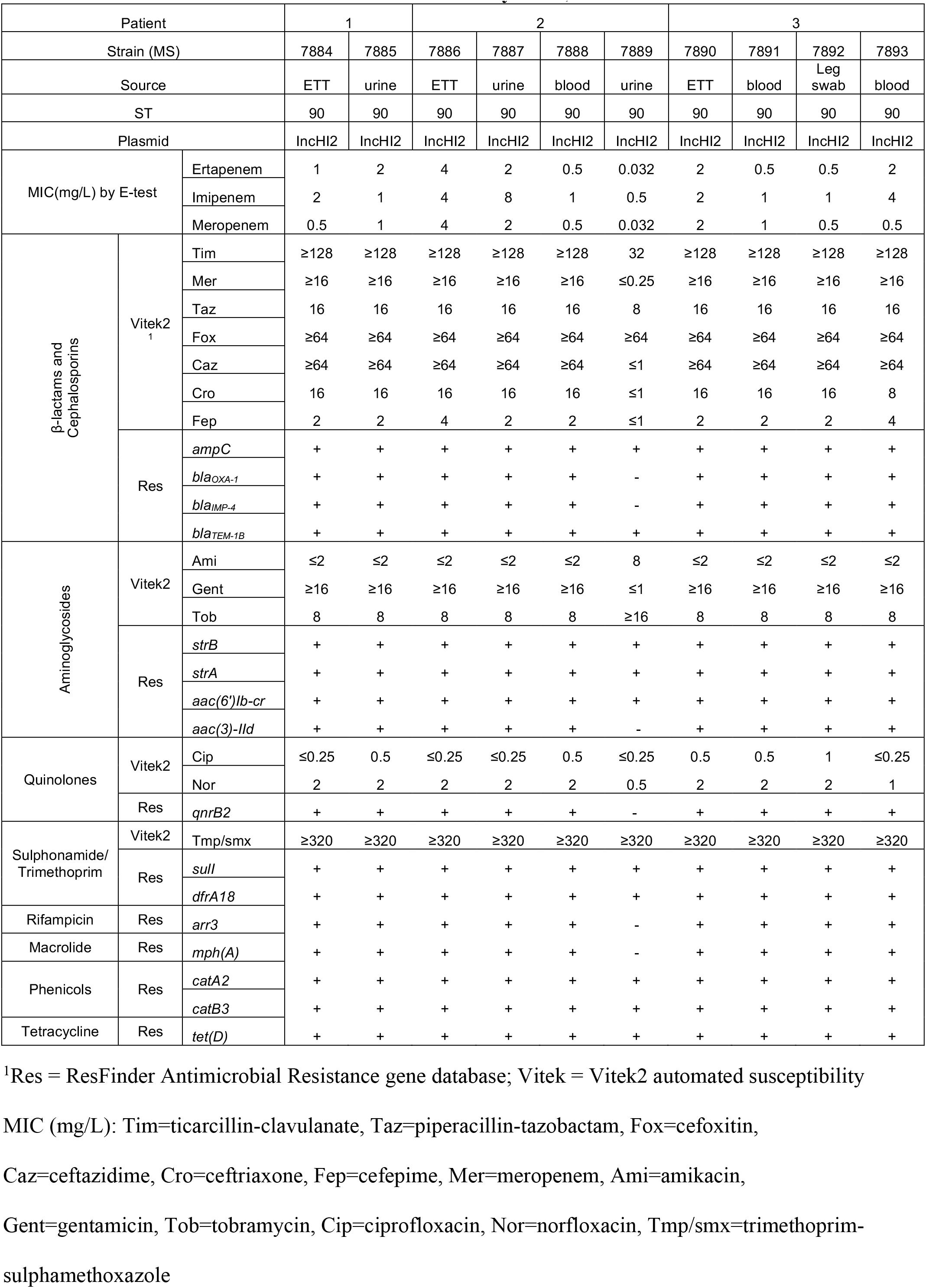
Antibiotic Resistance Profile as determined by Etest, Vitek2 and ResFinder

Patient 1 underwent debridement and split skin grafting for 29% total body surface area burns on day 2 of ICU admission and subsequently had 3 procedures in the burns operating rooms (Figure 1). An additional MDR-*E. cloacae* complex isolate was isolated from urine on day 21, eight days after discharge from the ICU. After no further colonisation of MDR-*E. cloacae* complex, Patient 1 was discharged from the hospital on day 38.

Patient 2 underwent multiple grafting and debridement procedures and was discharged from the ICU on day 17 (Figure 1). MDR-*E. cloacae* complex colonisation from the ETT and from urine was noted on day 8 and day 15, respectively. By day 19, the patient developed clinical signs of sepsis, with a phenotypically identical isolate identified in blood cultures and from a central venous line (CVL) tip culture. She received piperacillin/tazobactam 4.5 grams 8-hourly for 2 days, improved following line removal and did not receive further antibiotics for this episode. A subsequent *E. cloacae* complex was isolated from urine collected from a urinary catheter 17 days later demonstrated a different antibiogram with susceptibility to third generation cephalosporins, meropenem and gentamicin. She received 3 days of oral norfloxacin 400mg twice daily with microbiological resolution.

Patient 3, a 39-year old woman, was admitted with 66% total body surface area burns to the same ICU 5 weeks after Patient 1 and 2 were admitted and 20 days after they had been discharged from the ICU (Figure 1). MDR-*E. cloacae* complex was cultured from the ETT of Patient 3 on day 12 of ICU admission. She had frequent brief admissions to several hospitals since 2010 (never to ICU), and no MDR Gram-negative bacilli were identified in clinical or screening samples during previous admissions. MDR-*E. cloacae* complex with *Pseudomonas aeruginosa* were isolated from 8 skin swabs and an additional ETT aspirate. On days 19 and day 21, MDR-*E. cloacae* complex was isolated from blood cultures in the context of skin graft breakdown and signs of systemic inflammatory response syndrome (SIRS) with increasing inotrope requirements (Figure 1). *Streptococcus mitis* was cultured from blood on day 19. On day 36, her condition worsened with signs of SIRS. Transesophageal echocardiography demonstrated aortic and mitral valve lesions consistent with endocarditis. Pancytopenia developed, with a bone marrow aspirate and trephine suggestive of peripheral consumption. Multiple suspected cerebral, pulmonary, splenic and renal septic emboli were identified on imaging. She was palliated on day 47 of admission due to extensive cerebral emboli (Figure 1).

## Materials & Methods

### Study setting

Primary isolates were obtained from patients admitted to the Royal Brisbane & Women’s Hospital (RBWH), a tertiary referral hospital with 929 beds in South-East Queensland, Australia. Additional IMP-producing isolates, cultured from patients admitted to other hospitals in the metropolitan Brisbane area (referred to as Hospital A and B), were obtained from Pathology Queensland – Central Microbiology for comparison (Table S4).

### Antimicrobial susceptibility testing and carbapenemase detection

All bacterial isolates were identified by matrix-assisted laser desorption/ionization mass spectrometry (MALDI-TOF) (Vitek MS; bioMérieux, France). Antimicrobial susceptibility testing was carried out using Vitek 2 automated AST-N426 card (bioMérieux) with Etest to determine MICs for meropenem, imipenem and ertapenem. Carbapenemase activity was assessed by the use of the Carba-NP test (RAPIDEC; bioMérieux) and the presence of the *bla*_IMP-4_-like carbapenemase gene confirmed using an in-house multiplex real-time PCR (also targeting NDM, KPC, VIM and OXA-48-like carbapenemases)^17^.

### Bacterial DNA extraction

Single colonies were selected from primary bacterial cultures and grown in 10 mL Luria Bertani (LB) broth at 37°C overnight (shaking 250 rpm). DNA was extracted using the UltraClean® Microbial DNA Isolation Kit (MO BIO Laboratories) as per manufacturer instructions.

### Genome sequencing, Quality Control and *de novo* Assembly

All isolates in this study were sequenced using Illumina (see supplementary appendix). Reads passing quality control (QC) were assembled using Spades v3.6.0^18^ under default parameters (without careful flag). Contigs with coverage less than 10x were removed from final assemblies. Final assembly metrics were checked using QUAST v2.3^19^ (Table S3). A single isolate (MS14449) was sequenced using Nanopore MinION sequencing (see supplementary appendix).

### Taxonomic identification

Illumina raw reads were analysed using Kraken v0.10.5-beta to determine species and possible contamination. With the exception of MS7925, which was found to be *E. coli*, initial analysis determined the isolates to be *E. cloacae.* Subsequent analysis of the *E. cloacae de novo* assemblies using MASH v2.0^20^ and FastANI v1.1^21^ against representative *E. cloacae* complex complete genomes (see supplementary material) determined all isolates to be *E. hormaechei*.

### Phylogenetic analysis

SHRiMP v2.2.3^22^ as implemented in Nesoni v0.130^23^ under default settings was used to determine core single nucleotide polymorphisms (SNPs) between the ten 2015 RBWH *E. hormaechei* genomes to the reference Ecl1 and create a minimal-spanning tree. Further details of the Ecl1 assembly and SNP-calling process are provided in the supplementary appendix. Maximum likelihood trees of Ecl1 and the 6 *E. cloacae* from Hospitals A and B were built using RAxML v8.1.15^24^ based on the Nesoni core SNPs. RAxML was run with the GTRGAMMA nucleotide substitution rate and an initial seed length of 456 (bootstrap 1000 with Lewis ascertainment correction). Core genome size was estimated using Parsnp v1.2^25^.

### Multi-locus Sequence Typing (MLST), Plasmid Typing and Antimicrobial Resistance (AMR) Gene Profiling

MLST of isolate raw reads was performed using srst2 v0.1.5^26^ with typing schemes available on PubMLST (http://pubmlst.org/). Plasmid replicon typing was done based on Compain *et al.*^27^. Antibiotic resistance genes were detected using the ResFinder database^28^ and the ARG-ANNOT database^29^ with BLASTn and srst2^26^ respectively. Manual confirmation was carried out using BLASTn and read mapping using Burrows-Wheeler Aligner (BWA v0.7.5a-r405)^30^. Further details of whole genome comparisons and phage analysis are given in the supplementary appendix.

### Pacific Biosciences (PacBio) Single Molecule Real-Time (SMRT) Sequencing

A representative *E. hormaechei* isolate from patient 1 (MS7884) was grown on LB agar at 37°C overnight. IMP positive colonies (determined by colony PCR) were grown overnight in 15 mL LB broth with 2 μg/mL meropenem to avoid plasmid loss. Genomic DNA was extracted using UltraClean® Microbial DNA Isolation Kit (MO BIO) as per manufacturer’s instructions. 18.7 μg of DNA was prepared for sequencing using an 8-12 kb insert library and sequenced on a PacBio RSII sequencer using 1 SMRT cell. Further details of the assembly, annotation methods and plasmid stability in MS7884 are given in the supplementary appendix.

### Metagenomic sequencing and analysis of environmental samples

Swab and water samples from the ICU and Burns Ward were collected July 2018. DNA was extracted directly from samples using the Qiagen DNeasy Powersoil extraction kit including Biospec Products 0.1 mm diameter glass beads (as per manufacturers instructions). Water samples were concentrated using a Whatman Nuclepore 0.2 μm polycarbonate 25 mm diameter filter prior to DNA extraction. Sample DNA was sequenced at the Australian Centre for Ecogenomics on an Illumina NextSeq 500 (see supplementary appendix).

All samples were screened for species using Kraken v1.0^31^. Samples were also screen for resistance genes using srst2 v0.2.0^26^ against the ARG-ANNOT database. MinHash sketches of the ST90 *E. hormaechei* chromosome MS7884 (GenBank: CP022532.1) and the associated IncHI2 plasmid pMS7884A (GenBank: CP022533.1) were generated using MASH v1.1.1^20^ at default settings. Illumina reads for each sample were screened against our reference sketches using the screen function in MASH. Samples that shared >=90% of hashes were mapped to the reference sequences. Mapped reads were then parsed using a custom script and *de novo* assembled using Spades v3.11.1 for MLST analysis using Abricate v0.8 (https://github.com/tseemann/abricate) and nucleotide comparison using ACT^32^ and BRIG^33^.

## Accession numbers

Genome data has been deposited under Bioproject PRJNA383436. Illumina raw reads (SRX2999336-SRX2999345, SRX5578807-SRX5578812, SRX5578814), Nanopore raw reads (SRX5578813), PacBio raw reads (SRX2999346-SRX2999347) and metagenomic reads (SRX5590605-SRX5590610) have been deposited in the Sequence Read Archive (SRA). The MS7884 chromosome (CP022532), pMS7884A plasmid (CP022533), and pMS7884B plasmid (CP022534) have been deposited in GenBank.

## Results

### All three patients carry carbapenemase-producing *E. cloacae* complex

With the exception of MS7889 (isolated from the urine of Patient 2 on day 36), all *E. cloacae* complex isolates collected from the outbreak were resistant to ceftriaxone, ceftazidime, ticarcillin-clavulanate, piperacillin-tazobactam, meropenem, gentamicin and trimethoprim-sulphamethoxazole by Vitek 2 testing (Table 1) and demonstrated carbapenemase production by Carba-NP. The MICs for meropenem were considerably lower when tested by Etest^34^, often falling below the clinically susceptible breakpoint defined by EUCAST, but above the epidemiological cut-off (ECOFF)^35^. MS7889 was fully susceptible to carbapenems (meropenem MIC=0.032 by Etest) and was negative for IMP-4-like genes by PCR (Table 1).

### Whole genome sequencing identifies a link to a previous IMP-producing isolate

WGS of 10 isolates from patients 1, 2 and 3 was initiated after an additional microbiological confirmation of a *bla*_IMP-4_ *E. cloacae* complex isolate from a third patient from the RBWH ICU (Figure 1). *In silico* analysis determined all to be sequence type (ST)90 *E. hormaechei* (part of the *E. cloacae* complex), with the majority exhibiting the same resistance gene profile, including a 100% identical *bla*_IMP-4_ gene (Table 1). The exception was the carbapenem susceptible isolate MS7889, which was confirmed by WGS to have lost the *bla*_IMP-4_ gene as well as several additional resistance genes conserved in the other *E. hormaechei* isolates (Table 1). All ten isolates contained an IncHI2 plasmid. Sequence analysis suggested that AmpC derepression was unlikely to contribute to carbapenemase activity in these strains (further details are given in the supplementary appendix).

Comparison of the *E. hormaechei* genomes to publicly available draft assemblies identified a close match to *E. hormaechei* Ecl1 (GenBank: JRFQ01000000; formerly *E. cloacae*), an ST90 strain isolated from a burns patient at the RBWH ICU almost two years prior to the 2015 outbreak^13,36^. Antibiotic resistance profiling of the Ecl1 genome revealed an identical resistance profile compared to the majority of the 2015 isolates (Table 1).

### The 2015 outbreak isolates were near identical at the core genome level to an isolate from 2013

To investigate the relationship between the isolates at single-nucleotide resolution, reads from the 2015 RBWH isolates were mapped to *E. hormaechei* draft assembly for Ecl1. All 2015 RBWH isolates differed by fewer than five core SNPs (4,934,357 bp core genome), consistent with a direct ancestral relationship (Figure 2). Two isolates from Patient 1 and two isolates from Patient 3 were indistinguishable at the core genome level (Figure 2), although all of the isolates from Patient 3 had lost a prophage region (refer supplementary appendix). Ecl1 (isolated in 2013) was very closely related to these isolates, differing by only one core SNP. All four isolates from Patient 2 contained a discriminatory single-nucleotide deletion, thereby ruling out Patient 2 to Patient 3 transmission (Figure 2).

**Figure 2:**
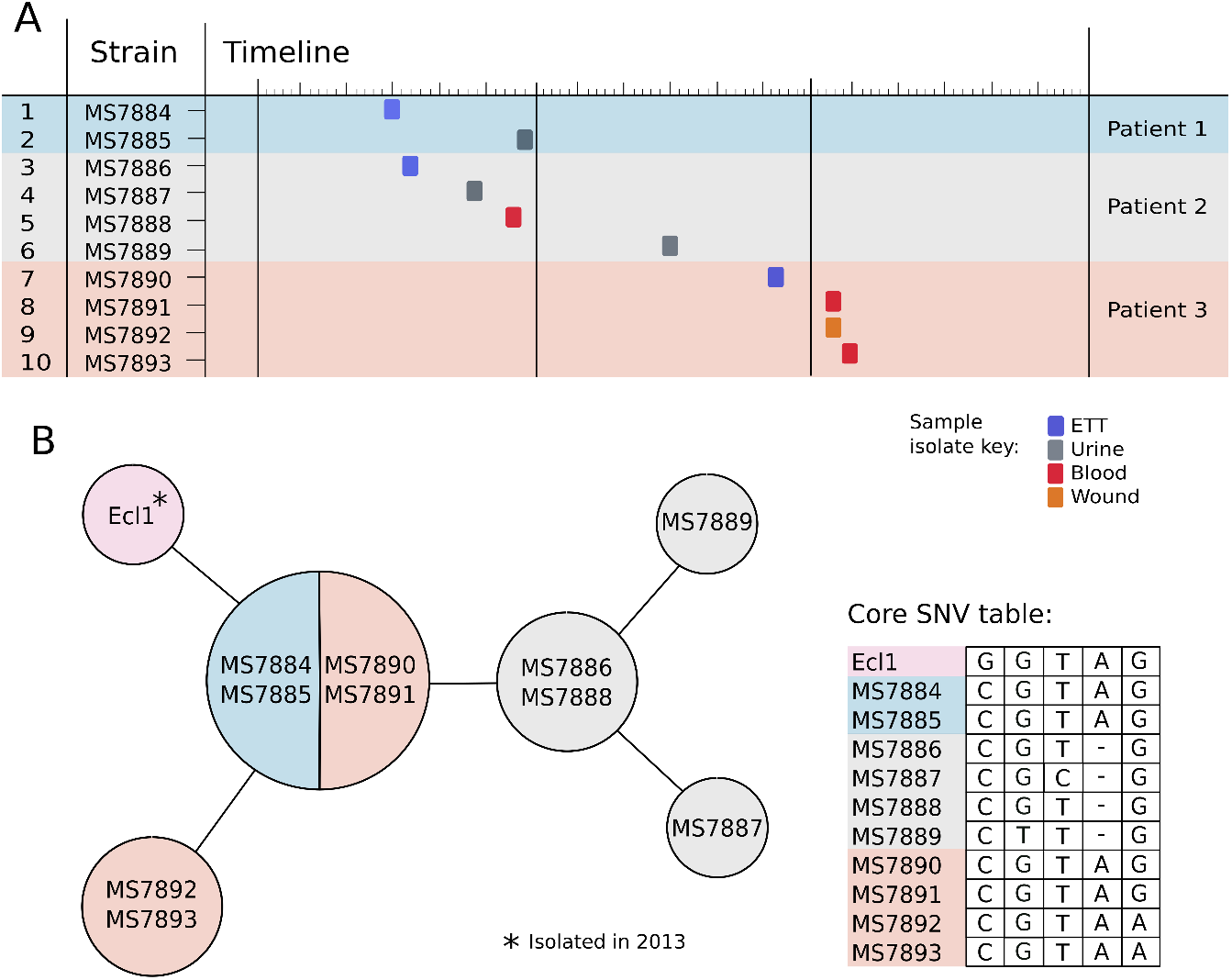
CPE isolate timeline and relationship matrix. **A**. 10 isolates were collected from 3 patients at various time-points in mid 2015. Coloured blocks indicate the source of the isolated strain: purple: respiratory, grey: urine, red: blood, and orange: wound. **B**. Relationship matrix (left) shows specific core single nucleotide variant (SNV) differences identified between strains. Strains within the same circle have identical core SNV profiles. Lines connecting circles represent accumulating SNV differences between strains (not-to-scale), where each line represents one SNV (including nucleotide deletion). Specific nucleotide differences between isolates are given in the table in panel B. Locations and consequences of nucleotide change are shown in Supplementary Dataset S1. All 11 isolates differed by 5 SNVs overall.

### Integration of WGS with infection control response

WGS analysis unequivocally linked all 10 isolates to the 2013 isolate Ecl1 from the same ward, confirming that the clone had not been an incursion from the accident affecting Patient 1 and 2 and that the hospital environment was suspected as the most likely original source of infection in the 2015 cases. In response, 28 environmental samples from the ICU, burns wards and operating theatres were collected 65 days after patient 1 and 2 were admitted and inoculated onto MacConkey agar with 8 mg/mL gentamicin (laboratory standard screening medium for MDR Gram-negative bacilli). No carbapenemase-producing *Enterobacter* spp. were detected. Additionally, no carbapenemase-producing *Enterobacter* spp. were detected in patients admitted to the ICU or burns unit for a 6-month period following the outbreak.

### Sequencing of additional CPE isolates identify a circulating IMP-4-carrying plasmid in Queensland

To determine the broader context of IMP-producing Enterobacteriaceae in surrounding hospitals, seven additional *bla*_IMP-4_ producing Enterobacteriaceae (*E. cloacae* complex n=6, *E. coli* n=1) were sequenced. These represented a selection of *bla*_IMP-4_ producing Enterobacteriaceae identified from Brisbane public hospitals via Pathology Queensland Central Microbiology for 2015. Both MLST and SNP analysis found no relationship to the 2015 RBWH *E. hormaechei*, with approximately 50,000 SNP differences between the ST90 representative strain Ecl1 and its nearest non-ST90 phylogenetic neighbour (Figure 3, also see supplementary appendix). Despite not being clonally related, all additional Enterobacteriaceae isolates possessed very similar antibiotic resistance gene profiles (Table S4), suggesting the possibility of lateral gene transfer via mobile genetic elements (e.g. integrons and/or plasmids). WGS analysis revealed that all 18 CPE isolates in this study, including the *E. coli* isolate, harbored an IncHI2 plasmid (plasmid ST1) and an identical *bla*_IMP-4_ gene, strongly suggesting plasmid-mediated circulation of *bla*_IMP-4_ between Enterobacteriaceae in Brisbane hospitals.

**Figure 3:**
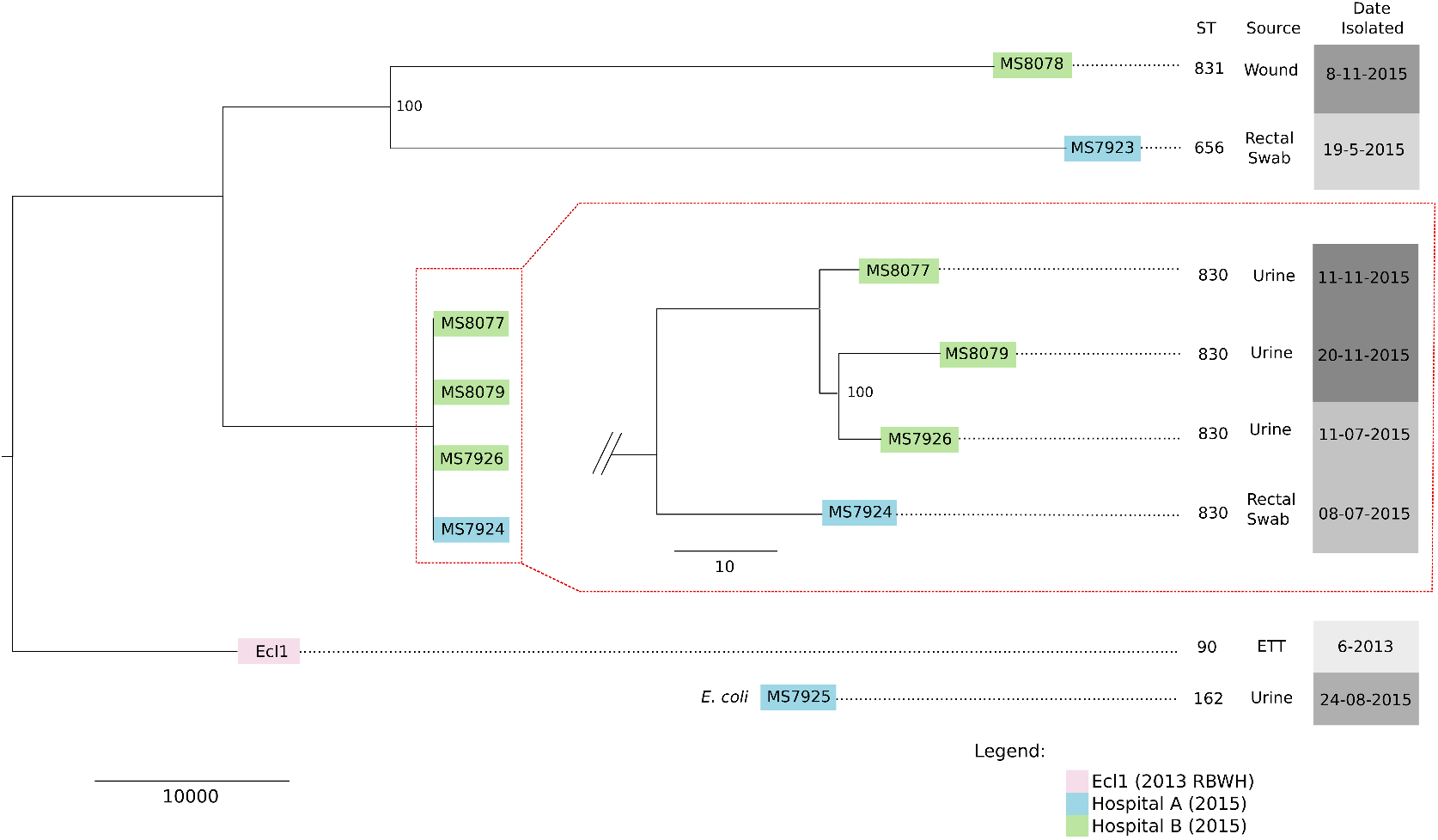
Core SNP Maximum likelihood (ML) tree of Hospital A and B *E. hormaechei* isolates in relation to RBWH isolates. Trimmed reads from 6 *E. hormaechei* isolates (Hospital A and B) were aligned to the reference *E. hormaechei* Ecl1 (isolated in 2013 at the RBWH) to determine core single nucleotide polymorphisms (SNPs) between all isolates. Ecl1 in this figure represents all 2015 RBWH isolates (n=10) as they were found to be near identical at the core genome level. 63,861 core SNPs were identified and used to generate a ML tree with RAxML (1000 bootstrap replicates), which determined no relationship between the RBWH isolates (pink) and the Hospital B (blue)/Hospital A (orange) isolates. Four closely related strains were identified from Hospitals A and B (red box). Alignment of trimmed reads from MS8077, MS8079 and MS7926 to MS7924 identified 117 core SNPs, however, a number of these SNPs were removed as they were identified as residing within transposon or phage regions. The remaining 58 core SNPs were used to generate a ML tree (1000 bootstrap replicates), showing that Hospital B strains differ by less than 20 SNPs.

### *bla*_IMP-4_ resides in the class 1 integron In809 on an IncHI2 plasmid

Due to the presence of multiple repetitive elements surrounding *bla*_IMP-4_, including insertion sequences (IS) and two suspected integrons with similar gene content, we were unable to accurately resolve the context of *bla*_IMP-4_ using Illumina sequencing alone. One representative isolate (MS7884) was sequenced twice using PacBio SMRT sequencing, which was able to resolve a complete closed chromosome of 4,810,853 bp and two plasmids: pMS7884A, a 330,060 bp IncHI2 plasmid carrying *bla*_IMP-4_ within a ~55 kb MDR region (Figure 4A), and pMS7884B, a smaller untypeable plasmid of 126,208 bp. The pMS7884A MDR region harbours two different class 1 integrons (In37 and In809) as well as a composite transposon conferring resistance to tetracycline and chloramphenicol (Figure 4A). BLASTn and read-mapping analysis revealed the presence of identical plasmids in all but one of the 18 isolates sequenced by Illumina in this study: isolate MS7889 is predicted to have lost a ~34 kb region from its MDR plasmid, including *bla*_IMP-4_, due to homologous recombination between two almost identical aminoglycoside resistance genes (Figure 4B). Notably in 15% of cases, sub-culture of MS7884 in the absence of meropenem selection resulted in loss of *bla*_IMP-4_ or the entire plasmid. Further details of the complete MS7884 genome and plasmid analysis are presented in the supplementary appendix.

**Figure 4:**
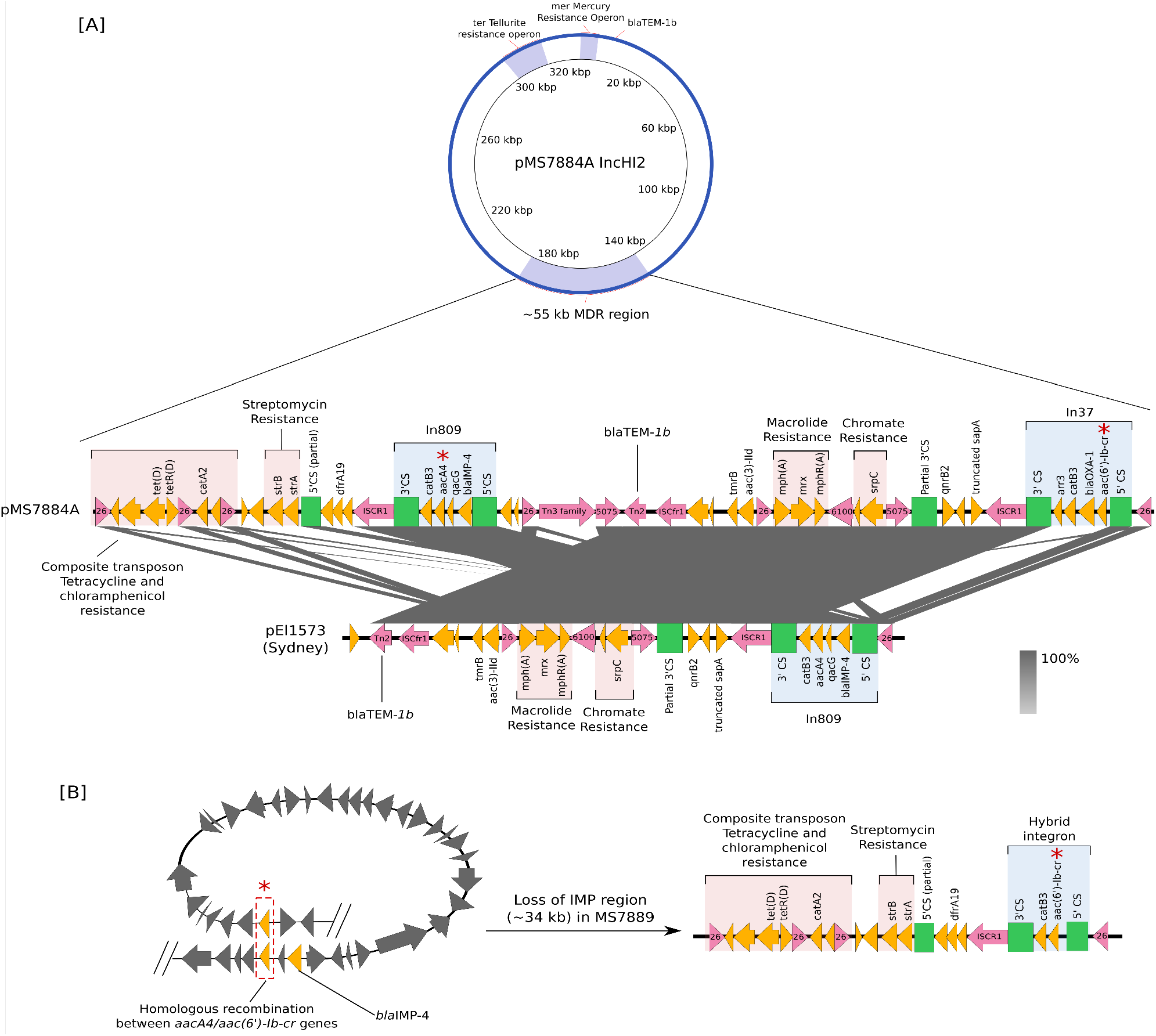
Large IncHI2 plasmid with ~55 kb multidrug resistance region containing IMP-4 carbapenemase. **A**. A 330,060 bp IncHI2 plasmid carrying multiple resistance operons, including a large ~55 kb multidrug resistance (MDR) region, was fully recovered and assembled using Pacific Biosciences (PacBio) SMRT sequencing of strain MS7884 (patient 1, isolate 1). The multidrug resistance region was found to contain two class 1 integrons (In809, In37) along with several other antibiotic resistance genes, as indicated. Comparison of this MDR region to publicly available genomes found a close match to pEl1573, isolated in 2012 from an *E. cloacae* isolate in Sydney, Australia. **B**. A predicted model of homologous recombination between two nearly identical *aacA4/aac(6’)-Ib-cr* genes (red asterisks) within the ~55 kb MDR region in MS7889 (patient 2, isolate 4, IMP-, carbapenem-susceptible) leading to the loss of a ~34 kb region containing *bla*_IMP-4_ as well as several other antibiotic resistance genes.

### Continued WGS surveillance reveals persistence of outbreak and transfer of *bla*_IMP-4_ plasmid

Ongoing surveillance has been in place for *bla*_IMP-4_ positive *E. hormaechei* isolates within the hospital since 2015. In 2016, during an unrelated outbreak, a *bla*_IMP-4_ positive *E. hormaechei* was isolated (MS14389) and following Illumina sequencing found to be only 2 SNPs different from the 2015 isolates, demonstrating continued persistence in the hospital environment (Figure S6).

In 2017, another *bla*_IMP-4_ positive *E. hormaechei* (MS14449) was isolated from the Hematology ward. Oxford Nanopore MinION sequencing has advanced in recent years to provide long-read sequencing and real-time analysis of bacterial isolates. However, no method has been established for its use in rapidly contextualising new isolates during ongoing outbreaks. Using the *de novo* Nanopore assembly and contextualising it against publicly available *E. cloacae* complex strains plus the outbreak isolates using whole genome alignment we were able to determine that this *E. hormaechei* was clonally unrelated to the 2015 outbreak isolates, but did carry an identical IncHI2 plasmid (Figure S7-S9). A *bla*_IMP-4_ positive *K. pneumoniae* isolate (MS14448) taken from the same patient was also sequenced (using only Illumina short-read sequencing) and found to carry a near identical IncHI2 plasmid, albeit missing a small section of the MDR region (Figure S10).

### Shotgun metagenomic sequencing reveals *E. hormaechei* ST90 and IncHI2 plasmid in hospital environment

Despite continued routine surveillance of the hospital environment using traditional culture methods an environmental source for the ST90 *E. hormaechei* was not found. In July 2018, 50 swab and water samples from the ICU and Burns ward environments were collected in response to an unrelated outbreak and subjected to both shotgun metagenomic sequencing and traditional culturing (Supplementary Dataset S2). From this round of surveillance, *Klebsiella oxytoca, E. cloacae* complex and *Leclercia adecarboxylata* were detected via traditional culturing methods from four samples (Table S7), however Illumina sequencing of these isolates determined that they were unrelated to the outbreak. Despite being clonally unrelated, three samples were found to carry a *bla*_IMP-4_-like gene (based on Real-Time PCR), which upon further inspection of the sequencing data corresponded to an IncHI2 plasmid with high similarity to pMS7884A (Supplementary Figure S11). While traditional culturing was unable to detect the ST90 *E. hormaechei*, metagenomic sequencing identified two samples with high confidence matches to the ST90 *E. hormaechei* reference MS7884 and the IncHI2 plasmid pMS7884A. Nucleotide comparison of the metagenomic assembled genomes (MAGs) for these samples (R5514 and R5537, both taken from floor drains) to our *E. hormachei* reference genome revealed a high level of nucleotide identity across the entirety of the chromosome and plasmid (Supplementary Figures 12 and 13). MLST analysis of both MAGs was also able to detect several similar, albeit incomplete, alleles for ST90 (Table S8). Additionally, screening for resistance genes identified the *bla*_IMP-4_ gene in both samples, further supporting the presence of an ST90 *E. hormaechei* and an IncHI2 plasmid similar to pMS7884A in these samples. Analysis of three other environmental samples (R5505, R5506 and R5522) recovered fewer than 5% of total reads that mapped to the reference *E. hormaechei* MS7884 genome, compared to 25% and 20% for our positive samples R5514 and R5537 respectively (Table S8, Supplementary dataset 2). This low number of reads was insufficient to confidently identify the ST90 *E. hormaechei* or IncHI2 plasmid of interest in these samples (Table S8 and Supplementary Figures 12 and 13).

## Discussion

While there has been a dramatic improvement in the cost and availability of whole genome sequencing (WGS), it is not clear how these advances can best be incorporated into routine clinical microbiology. Several studies have demonstrated the ability of WGS to provide optimal discrimination between strains to help inform a response to outbreaks or nosocomial acquisition^37–40^. Here, we demonstrate that and integrated WGS approach can help rapidly characterize an outbreak in a critical care setting, particularly regarding transmission pathways. We highlight how WGS can be used to link contemporary outbreak isolates to historical isolates to inform infection control and incorporate long-read sequencing technologies to resolve complete genomes (including plasmids) and to rapidly resolve suspected outbreak cases. Finally, we demonstrate the potential of using complete genome sequences to interrogate environmental shotgun metagenomic sequencing data to identify outbreak sources.

The finding that the outbreak strains were virtually indistinguishable from an IMP-4-producing *E. hormaechei* isolated two years previously from the same unit was unexpected and highlighted the need to consider environmental sources and potential person-to-person transmission, as has been previously described in Australian ICU and burns units^14^. Despite ongoing surveillance, traditional culture-based detection methods were unable to find the ST90 *E. hormaechei* in the environment. Direct DNA extraction and metagenomic sequencing has in recent years revolutionised infectious disease surveillance, allowing detection of all microorganisms in a sample without the biases and limitations of traditional pathogen detection^41^. Using metagenomic sequencing and a high quality complete reference genome we were able to detect two samples with high confidence hits to an ST90 *E. hormaechei* and an IncHI2 plasmid, confirming its presence in the environment. We were also able to observe the overall community profile within each environmental sample, making metagenomic sequencing a powerful infection control and surveillance tool for tracking (i) the types of bacteria present in the environment, (ii) the types of resistance genes circulating, and (iii) the effectiveness of environmental cleaning. Metagenomic sequencing does, however, have certain limitations when considering its implementation in routine infection control, including the necessity for a reasonable amount of starting DNA, and the chance of amplification inhibition during library preparation (causing low yield or failed sequencing). Metagenomic sequencing can also be quite costly^42^, as the sequencing output needs to be sufficiently high to provide an accurate population structure and detect low-abundance organisms. In our study, while metagenomic sequencing of the environmental samples yielded positive results, it is still unclear how these reservoirs are causing reinfection in patients. It is possible that healthcare workers are somehow involved, with previous studies confirming carriage of a range of clinically important bacteria^43–45^.

Using SMRT sequencing technology, we determined the full context of *bla*_IMP-4_ and its location within a large, complex and highly repetitive MDR region harbouring two integrons: In37 and In809. In37 is a widespread class 1 integron that has been found in many bacterial species^46,47^. In809, which carries *bla*_IMP-4_, has previously been described from *K. pneumoniae* (GenBank: KF250428.1, HQ419285.1, AJ609296.3), *E. cloacae* (GenBank: JX101693.1) and *Acinetobacter baumannii* (GenBank: AF445082.1, DQ532122.1) in various plasmid backgrounds including IncA/C2^48^, IncL/M and IncF^49^. Most recently, a carbapenemase-producing *Salmonella* sp. isolated from a domestic cat in Australia was shown to contain *bla*_IMP-4_ within an IncHI2 MDR plasmid (pIMP4-SEM1)^50^. Remarkably, we found that pIMP4-SEM1 was near identical to pMS7884A (Figure S5). This finding highlights the role of domestic animals (or the food they eat) as a reservoir for antibiotic resistance genes.

Analysis of several CPE in this study suggested that a common plasmid or integron carrying multiple antibiotic resistance genes is likely the major driver of antibiotic resistance dissemination across a broad range of Enterobacteriaceae. In addition to the presence of *bla*_IMP-4_, four resistance genes (blaTEM-1b, *bla*_IMP-4_, *qnrB*, and *aac(6’)-Ib*) carried by these isolates were previously detected by PCR in the majority of 29 IMP-4-producing *E. cloacae* complex isolates surveyed from Queensland hospitals between June 2009 to March 2014^13^. Only one of these isolates was ST90, suggesting lateral transfer of these genes to different *Enterobacter* clones in Queensland before 2013. During ongoing surveillance at RBWH we identified an *E. hormaechei* isolate that also carried the *bla*_IMP-4_ plasmid, but was not closely related to the outbreak isolates, suggesting ongoing lateral transfer within the hospital environment. In this instance the Oxford Nanopore MinION sequencing platform was instrumental in ruling out this isolate from the ongoing outbreak of *bla*_IMP-4_ ST90 *E. hormaechei*. Our work highlights the potential for integrating this highly portable and rapid technology as part of the “genomic toolkit” for infection control alongside more established platforms. The capacity to recognise inter-species MDR plasmid transfer is also greatly enhanced by the availability of complete reference genomes, as demonstrated by the *bla*_IMP-4_ positive *K. pneumoniae* isolate identified in this study.

There were significant discrepancies between meropenem MICs according to the testing modality used, with the Etest consistently testing as “susceptible/intermediate” (MIC ≤4 mg/L; range 0.5-4 mg/L) and Vitek2 as “resistant” (usually with MICs ≥16 mg/L). According to pharmacokinetic/pharmacodynamic (PK/PD) principles, provided the MIC to a carbapenem falls within a susceptible range, the agent may still be effective despite the presence of a carbapenemase^51^. Robust clinical data to help guide therapy are lacking and many clinicians rely on combination therapy to optimize efficacy against carbapenemase-producers, largely based on observational studies suggesting benefit^52,53^. The presence of carbapenemase genes may be missed if clinical breakpoints for carbapenem MICs are used^35^, however it can be rapidly ascertained by WGS, without *a priori* assumptions of which genes are likely to be present. A wealth of additional information that may influence clinical decisions can be obtained, such as the presence of other β-lactamases, factors that may regulate resistance gene expression (e.g. IS elements), mutations in outer-membrane proteins, or other known resistance genes.

## Conclusions

We used an integrated WGS approach to help elucidate genetic relationships between *bla*_IMP-4_ carbapenemase-producing *E. hormaechei* identified from our ICU and Burns facility. Real-time application of this technology revealed an unexpected clonal relationship with a strain isolated from the same unit two years previously. Continued routine WGS surveillance has enabled detailed monitoring of the outbreak, with rapid nanopore sequencing crucial for ruling out a suspected case in a previously unaffected ward. Comparisons with other Enterobacteriaceae containing *bla*_IMP-4_ isolated from surrounding hospitals revealed its carriage on a broad host range IncHI2 plasmid, assumed to be circulating via lateral gene transfer across different *E. cloacae* complex clones and also *E. coli.* SMRT sequencing enabled the genetic context of all resistance genes within this plasmid to be resolved and revealed the mechanism of loss of resistance genes in one *E. hormaechei* strain that reverted to a fully carbapenem-susceptible phenotype. The availability of a complete *E. hormaechei* reference chromosome and *bla*_IMP-4_ plasmid were also instrumental in locating a suspected source of the outbreak in the hospital plumbing with shotgun metagenome sequencing. As WGS technologies become increasingly available, they are likely to prove essential tools for the clinical microbiology laboratory to respond to emergent infection control threats, and can be used in real-time to provide clinically meaningful information.

## Acknowledgements

We thank Krispin Hajkowicz, Trish Hurst and Michelle Doidge (Infectious Diseases Department, RBWH, Brisbane, Australia) for collecting environmental samples used in this study.

