## Supplementary Appendix for "Integrating multiple genomic technologies to investigate an outbreak of carbapenemase-producing *Enterobacter hormaechei*"

1. School of Chemistry and Molecular Biosciences, The University of Queensland, Brisbane, QLD, Australia; 2. Australian Infectious Diseases Research Centre, The University of Queensland, Brisbane, QLD, Australia; 3. Australian Centre for Ecogenomics, The University of Queensland, Brisbane, QLD, Australia; 4. The University of Queensland, UQ Centre for Clinical Research, Brisbane, QLD, Australia; 5. Pathology Queensland, Central Microbiology, Brisbane, QLD, Australia; 6. Burns Trauma and Critical Care Research Centre, The University of Queensland, Brisbane, QLD, Australia; 7. Infectious Disease Unit, Royal Brisbane & Women's Hospital, Brisbane, QLD, Australia; 8. Division of Genetics and Molecular Biology, Institute of Biological Sciences, Faculty of Science, University of Malaya, Kuala Lumpur, Malaysia; 9. Wesley Medical Research, The Wesley Hospital, Auchenflower, QLD, Australia

#### Appendix contents:

**Supplementary Methods** 3

*Illumina Sequencing and Quality Control (QC)* 3
*Taxonomic identification* 4
*Determining reference for phylogenetic analysis* 5
*Reassembly of Ecl1 from raw reads* 5
*Whole genome comparisons and phage analysis* 6
*Assembly and Annotation of SMRT sequenced isolate MS7884* 6
*Nanopore MinION sequencing of isolate MS14449* 7
*Colony polymerase chain reaction (PCR) to identify plasmids in MS7884* 8

**Supplementary Results** 9

*Table S3: Assembly metrics* 9
*Table S4: Hospital A and B isolates Resistance gene profile* 10
*Phenotypic observations may relate to single nucleotide polymorphisms (SNPs)* 11
*Patient 3 isolates are missing ~25 kb phage region* 11
*AmpC is unlikely to contribute to carbapenem resistance in the 2015 RBWH isolates* 12
*Loss of ~34 kb region in MS7889 is mediated by homologous recombination* 13
*Unique SNPs in aac(6')-Ib-cr could result in increased resistance in MS7889* 13
*Whole genome sequencing identifies two plasmids carried in RBWH isolates* 14
*pMS7884A and pMS7884B comparisons to published genomes* 15
*Summary of methylomes in complete PacBio genomes* 15
Table S5: Summary of recognition motifs[1] in MS7884A and MS7884B 16
Table S6: Summary of methyltransferases 17
*Continued surveillance using Nanopore MinION sequencing* 17
*Environmental surveillance in the hospital* 18
Table S7: Isolates cultured from environmental surveillance swabs 18
Table S8: MLST alleles detected in MAGs from samples with *E. hormaechei* detected based on MASH 19

**References** 20

**Supplementary Figures** 22

- *Figure S1: Whole genome comparison of RBWH draft assemblies using BRIG*
- *Figure S2: Nucleotide alignment between MS7884 aac(6')-Ib-cr, MS7884 aacA4 and MS7889 aac(6')-Ib-cr gene*
- *Figure S3: Multiple sequence alignment of aacA4 and aac(6')-Ib-cr amino acid sequence in MS7884 and MS7889*
- *Figure S4: BRIG comparison of 2015 RBWH draft assemblies to pECL_A from E. cloacae subsp. cloacae ATCC 13047*
- *Figure S5: Easyfig comparison of pIMP4-SEM1 and pMS7884A*
- *Figure S6: Relationship matrix of outbreak ST90 E. hormaechei with MS14389*
- *Figure S7: Comparison of large plasmid from MS14449 and pMS7884A*
- *Figure S8: BRIG comparison of pMS7884B and small plasmid from MS14449*
- *Figure S9: Parsnp tree contextualising MS14449 Illumina and Nanopore draft assemblies*
- *Figure S10: BRIG comparison of MS14448 and MS14449 draft assemblies against pMS7884A*
- *Figure S11: BRIG comparison of isolate draft assemblies from metagenomic sequencing against pMS7884A*
- *Figure S12: BRIG of MAGs from samples with positive E. hormaechei*
- *Figure S13: BRIG of MAGs from samples with positive pMS7884A*

### Supplementary Methods

#### Illumina Sequencing and Quality Control (QC)

**<u>Original outbreak isolates (2015)</u>**

Illumina sequencing of the 2015 *bla*IMP+ *E. hormaechei* isolates (n=16) and *E. coli* isolate (n=1) was done at the Australian Centre for Ecogenomics (ACE). *E. hormaechei* Ecl1 (formerly identified as *E. cloacae*) had previously been sequenced on Illumina HiSeq2000 at the Australian Genome Research Facility (Melbourne, Australia). Libraries for isolates MS7884-MS7893 (RBWH isolates) were prepared using 2 x 150 bp Nextera XT V1 chemistry and run on an Illumina NextSeq. Isolates MS7923-MS7926 and MS8077-MS8079 (Hospitals A and B) were prepared using 2 x 300 bp Nextera XT libraries with V3 chemistry and run on an Illumina MiSeq. All libraries were quantified using Viaa7 Thermo Fisher qPCR with QC checks with bioanalyzer analysis (Agilent). Contaminant searches on the raw reads were performed using Kraken (v0.10.5-beta). All raw reads were filtered using Nesoni (v0.130) [1] to remove Illumina adaptor sequences, reads shorter than 80 bp and bases below Phred quality 5. MiSeq reads were additionally hard trimmed to 150 bp due to low quality bases between 1-10 and 160-300 using Nesoni (v0.130).

**<u>Sequencing of additional isolates at Queensland Forensic Scientific Services (QFSS)</u>**

All isolates collected subsequent to the original outbreak were sequenced at QFSS (MS14449, MS14448, MS14389, M87132, M87133, M87134, and M87135). DNA was extracted using the DSP DNA Mini Kit on the QIAsymphony SP (Qiagen). Libraries were prepared using the Nextera XT DNA preparation kit (Illumina) and the sequencing was performed on the NextSeq 500 (Illumina) with 2x150bp chemistry, NextSeq Midoutput kit v2.5.

**Metagenomic sequencing at Australian Centre for Ecogenomics**

*Library Preparation*

Libraries were prepared according to the manufacturer's protocol using Nextera XT Library Preparation Kit (Illumina #FC-131-1096). The only alterations to the protocol as outlined was the reduction of total reaction volume for processing in 96 well plate format. Library preparation and bead clean-up was run on the Mantis Liquid Handler (Formulatrix) and Epimotion (Eppendorf #5075000301) automated platform. These programs cover "Tagment Genomic DNA" to "Amplify DNA" in the protocol (Mantis- Nextera XT library prep protocol) and "Clean Up Libraries" in the protocol (Epimotion - Library Clean Up protocol). On completion of the library prep protocol, each library was quantified and QC was performed using the Qubit™ dsDNA HS Assay Kit (Invitrogen) and Agilent D5000 HS tapes (#5067-5592) on the TapeStation 4200 (Agilent # G2991AA) as per the manufacturer's protocol.

*Library Pooling, QC and Loading*

Nextera XT libraries were pooled at equimolar amounts of 1nM per library to create a sequencing pool. The library pool was quantified in triplicates using the Qubit™ dsDNA HS Assay Kit (Invitrogen). Library QC is performed using the Agilent D5000 HS tapes (#5067-5592) on the TapeStation 4200 (Agilent # G2991AA) as per the manufacturer's protocol. The library was prepared for sequencing on the NextSeq500 (Illumina) using NextSeq 500/550 High Output v2 2 x 150bp paired end chemistry in the Australian Centre for Ecogenomics according to manufacturer's protocol.

**Taxonomic identification**

*De novo* assemblies for all *E. cloacae* complex isolates were compared to representative publicly available complete *E. cloacae* complex genomes using MASH v2.0 [2] and FastANI v1.1 [3] at default settings. Genome comparisons were made against *Enterobacter asburiae* (NZ_CP011863.1), *Enterobacter bugandensis* (NZ_LT992502.1), *Enterobacter clocace* subsp.

*cloacae* (NC_014121.1), *Enterobacter hormaechei* subsp. *steigerwaltii* (NZ_CP017179.1), *Enterobacter kobei* (NZ_CP017181.1), *Enterobacter ludwigii* (NZ_CP017279.1), *Enterobacter roggenkampii* (NZ_CP017184.1), *Enterobacter hormaechei* subsp. *xiangfangensis* (NZ_CP017183.1), and *Enterobacter hormaechei* subsp. *hoffmannii* (NZ_CP017186.1). The top hit, based on %ANI and greatest number of matching hashes (for fastANI and MASH respectively), was used to identify the species for that isolate.

##### Determining reference for phylogenetic analysis

Kraken (v0.10.5-beta) analysis using raw reads from the ten 2015 RBWH strains determined *E. cloacae* subsp. cloacae NCTC 9394 (GenBank: FP929040.1) to be the closest match in the complete genome division of GenBank. Subsequent comparison of the ten 2015 RBWH draft genome assemblies to assemblies within the publicly available whole-genome shotgun (WGS) database revealed a closer match to the IMP-4-producing *E. hormaechei* strain Ecl1 from 2013 [4] (Accession: JRFQ01000001; formerly identified as *E. cloacae*). Both were subsequently used to confirm single nucleotide variations (SNVs) between the 2015 RBWH strains.

##### Reassembly of Ecl1 from raw reads

In order to determine the relationship between *E. hormaechei* Ecl1 and the 2015 RBWH *E. hormaechei* strains, core SNP distances were determined using a mapping approach as implemented through Nesoni v0.130 [1] against the publicly available Ecl1 genome (Accession: JRFQ01000001). Manual inspection of the Nesoni output identified a number of clustered SNPs that we suspected were erroneous (Supplementary dataset S1). Reassembly of the raw reads using Spades v3.6.0 [5] under default parameters (without careful flag) was unable to correct the erroneous SNPs (Supplementary dataset S1). Closer examination of the read pileups for Ecl1 identified strand-specific clusters of SNPs, possibly due to technical problems at the time of sequencing. To identify genuine variants, the read pileup for every SNP as reported by Nesoni was manually curated using Artemis and Bamview. SNPs that were due to strand-specific base call errors in the assembly, resided close to contig edges, or resided in repetitive region (e.g. insertion sequences) were omitted from downstream analyses.

##### Whole genome comparisons and phage analysis

Whole genome comparisons were performed using BRIG [6]. Regions of difference were further investigated using the Artemis Comparison Tool (ACT) [7]. A ~25 kb region, missing in all patient 3 strains, was analysed using BLASTn to determine predicted gene content. As the region was predicted to contain phage-like proteins, Ecl1, MS7884 (RBWH patient 1), MS7886 (RBWH patient 2) and MS7890 (RBWH patient 3) draft assemblies were analysed using PHAST [8] to further characterise the phage-like region.

##### Assembly and Annotation of SMRT sequenced isolate MS7884

MS7884 was first sequenced on a PacBio RSII sequencer using the P6-C4 sequencing chemistry at the University of Malaya and assembled using the SMRT Analysis suite (version 2.3.0) to give a single chromosome (MS7884B: 4,810,853 bp) and one un-typeable plasmid (pMS7884B: 126,208 bp). However, neither pMS7884B nor the chromosome contained any of the previously identified antibiotic resistance genes, including the carbapenemase *bla*$_{IMP-4}$ (previously identified using Illumina and PCR). To ensure that the suspected multidrug resistant (MDR) plasmid was retained, we re-extracted DNA from MS7884 grown with 2 µg/mL meropenem (in lysogeny broth, LB). Sequencing was carried out using a different PacBio RSII Sequencer (P6-C4 sequencing chemistry) at the Doherty Institute, University of Melbourne. Assembly of this genome using the SMRT Analysis suite (version 2.3.0) gave an identical chromosome (4,810,853 bp) and a single IncHI2 plasmid (330,060 bp) carrying *bla*$_{IMP-4}$

(hereafter referred to as pMS7884A). To avoid duplication in the complete genome databases we have compiled the MS7884A chromosome, pMS7884A plasmid and pMS7884B plasmid as a single strain MS7884 genome, consistent with our original observation (according to Illumina sequencing) that both plasmids were present in the original patient isolate. Individual PacBio Raw data files are available to enable comparisons of the methylome between the two spontaneously cured MS7884 derivative isolates (see 'Summary of methylomes in complete PacBio genomes' below).

Table S1: Complete chromosome and plasmid list

| Lab number | Parent strain | Chromosome/ plasmid | Size (bp) | Isolate name | Feature |
|---|---|---|---|---|---|
| MS7969 | MS7884 (single colony) | Chromosome | 4,810,853 | MS7884B | |
| | | Plasmid | 126,208 | pMS7884B | No IMP carbapenemase gene |
| MS8407 | MS7884 (single colony grown with 2 µg/ml meropenem) | Chromosome | 4,810,853 | MS7884A | |
| | | Plasmid | 330,060 | pMS7884A | Contains multiple antibiotic resistance genes, including IMP-4 carbapenemase |

All PacBio SMRT sequences were manually closed using the Artemis Comparison Tool (ACT) [7]. The chromosome and plasmids were polished using both the PacBio reads (with Quiver through the SMRT Analysis suite) and the Illumina reads (using BWA v0.7.5a-r405 with consensus calling through Pilon v1.21) to remove erroneous indels. The final genome was annotated using Prokka (v1.12-beta). The integron regions were manually annotated using a combination of RAC [9], as implemented through MARA (http://app.spokade.com/mara/), and Integrall [10]. Insertion sequences (IS) were manually annotated using ISSaga [11]. Modified bases and associated motifs were detected using the SMRT analysis suite (version 2.3.0) to determine genome-wide methylation.

#### Nanopore MinION sequencing of isolate MS14449

Hospital surveillance identified a *bla*$_{IMP-4}$ positive *E. hormaechei* isolate in October 2017 from a patient in the Hematology ward of RBWH (MS14449). MS14449 was grown overnight on horse blood agar at 37°C. DNA was extracted using the MoBio UltraClean Microbial DNA isolation kit (as per manufacturer's instructions). 1.5 ug of DNA was prepared using the 1D Genomic DNA by Ligation sequencing kit (SQK-LSK108 version GDE_9002_v108_revT_18Oct2016). The entire library was loaded onto a MIN106 R9.4 flow cell and run for 26 hours using MinKNOW version 1.7.14 on a Mac OSX operating system. A subset of 80,000 fast5 files were basecalled using Albacore (v1.1.1). Basecalled reads were filtered for length (minimum 2000 bp) and quality (minimum 10) using Japsa (https://github.com/mdcao/japsa). The remaining 62,093 reads were assembled using Canu (v1.3) at default settings resolving a single chromosome and two plasmids.

#### Colony polymerase chain reaction (PCR) to identify plasmids in MS7884

MS7884 was grown on LB agar overnight at 37°C. Single colonies were resuspended in 50 µL sterile $dH_20$, boiled for 15 minutes and centrifuged at 280x g for 2 minutes. 5 µL of supernatant containing colony DNA was used in 25 µL OneTaq PCR reactions, as per manufacturer's recommendations (New England Biolabs Inc.). PCR primers were designed to identify both plasmids (Table S2). PCR was run as follows: Annealing 30 seconds at 55°C, extension 1 minute at 68°C, denaturing 30 seconds at 94°C, 29 cycles.

**Table S2: Primer list**

| Plasmid Target | Primer Target | Product size | tm | Sequence |
|---|---|---|---|---|
| ~330 kb IncHI2 | MS7884_pA0187 BlaIMP-4 | 200 | 58 | AGGACACACTCCAGATAACC |
| | | | 67 | TGATGCGTCTCCAGCTTCAC |
| | MS7884_pA0259 RepHI2 | 800 | 58.88 | TAATGGAGAGCGAGGGGTTC |
| | | | 58.62 | GCGGTTAAATCATGGACGGT |
| ~126 kb un-typeable | MS7884_pB0001 Putative methyltransferase | 420 | 59.21 | GAAATGTACCGCGTGCTGAA |
| | | | 58.96 | TCCTCAAGCATCTCGATCCC |
| | MS7884_pB0009 Replication initiation protein repE | 590 | 58.42 | AGAATAGCCCGCGAATTGTC |
| | | | 60.04 | CAGGAACCTACGGCGAAAGT |

#### Supplementary Results

Table S3: Assembly metrics

| Strain | Location | Depth[2] | Contigs[3] | Largest contig | N50 | Total Length |
|---|---|---|---|---|---|---|
| Ecl1 (JRFQ01000001) | RBWH | 30x | 83 | 514 952 | 175 645 | 5 199 581 |
| Ecl1 (reassembly) [1] | RBWH | 36x | 90 | 515 066 | 256 013 | 5 223 643 |
| MS7884 | RBWH | 93x | 84 | 556 131 | 259 415 | 5 225 529 |
| MS7885 | RBWH | 126x | 87 | 515 066 | 208 661 | 5 226 022 |
| MS7886 | RBWH | 85x | 85 | 556 628 | 259 182 | 5 228 488 |
| MS7887 | RBWH | 104x | 86 | 385 936 | 259 128 | 5 214 118 |
| MS7888 | RBWH | 99x | 86 | 406 431 | 259 182 | 5 227 406 |
| MS7889 | RBWH | 80x | 79 | 357 241 | 195 846 | 5 181 005 |
| MS7890 | RBWH | 91x | 84 | 406 431 | 208 661 | 5 195 590 |
| MS7891 | RBWH | 124x | 83 | 556 346 | 283 853 | 5 195 194 |
| MS7892 | RBWH | 121x | 89 | 388 759 | 259 518 | 5 194 114 |
| MS7893 | RBWH | 140x | 83 | 515 066 | 259 182 | 5 193 144 |
| MS7923 | Hospital A | 109x | 61 | 899 556 | 301 561 | 5 007 334 |
| MS7924 | Hospital A | 76x | 73 | 1 033 606 | 327 391 | 5 150 594 |
| MS7925 | Hospital A | 91x | 96 | 622 503 | 143 352 | 5 137 892 |
| MS7926 | Hospital B | 98x | 83 | 721 578 | 282 470 | 5 203 172 |
| MS8077 | Hospital B | 98x | 82 | 588 399 | 307 036 | 5 209 475 |
| MS8078 | Hospital B | 71x | 116 | 329 971 | 146 076 | 5 028 820 |
| MS8079 | Hospital B | 103x | 83 | 733 216 | 261 436 | 5 161 736 |

[1] Genome reassembled to try and fix erroneous SNPs
[2] Average depth of coverage calculated through Nullarbor
[3] Number of contigs > 78 bp in length

#### Table S4: Hospital A and B isolates Resistance gene profile

| Strain | Species | Date | Location | Source | ST | Plasmid | Beta-lactam resistance | | | | | Rif | Aminoglycoside resistance | | | | Quinolone resistance | | Phe | | MLS | Sul | Trim | Tet | Fos |
|---|---|---|---|---|---|---|---|---|---|---|---|---|---|---|---|---|---|---|---|---|---|---|---|---|---|
| | | | | | | | *ampC* | $bla_{SHV\text{-}12}$ | $bla_{OXA\text{-}1}$ | $bla_{IMP\text{-}4}$ | $bla_{TEM\text{-}1B}$ | *arr3* | *strB* | *strA* | *aac(6')Ib-cr* | *aac(3)-IId* | *qnrA1* | *qnrB2* | *catA2* | *catB3* | *mph(A)* | *sul1* | *dfrA18* | *tet(D)* | *fosA* |
| MS7925 | *E. coli* | 17/06/15 | A | Urine | 162 | IncHI2 | + | + | + | + | + | + | + | + | + | + | - | + | + | + | + | + | + | + | - |
| MS8078 | *E. hormaechei* | 8/11/15 | B | Swab Wound | 831 | IncHI2 | + | + | + | + | + | + | + | + | + | + | - | + | + | + | + | + | + | + | + |
| MS7923 | *E. hormaechei* | 19/05/15 | A | Rectal swab | 656 | IncHI2 | + | - | + | + | + | + | + | + | + | + | - | + | + | + | + | + | + | + | + |
| MS8077 | *E. hormaechei* | 11/11/15 | B | Urine | 830 | IncHI2 | + | - | + | + | + | + | - | - | + | + | + | + | + | + | + | + | + | + | + |
| MS8079 | *E. hormaechei* | 20/11/15 | B | Urine | 830 | IncHI2 | + | - | + | + | + | + | - | - | + | + | - | + | + | + | + | + | + | + | + |
| MS7926 | *E. hormaechei* | 11/07/15 | B | Urine | 830 | IncHI2 | + | - | + | + | + | + | - | - | + | + | + | + | + | + | + | + | + | + | + |
| MS7924 | *E. hormaechei* | 8/07/15 | A | Rectal swab | 830 | IncHI2 | + | - | - | + | + | + | - | - | + | - | - | + | + | + | + | + | + | + | + |
| MS7884 [1] | *E. hormaechei* | 15/06/15 | RBWH | ETT | 90 | IncHI2 | + | - | + | + | + | + | + | + | + | + | - | + | + | + | + | + | + | + | - |

Abbreviations: Rif = rifampicin, Tet = Tetracycline, Trim = Trimethoprim, Sul = Sulphonamides, MLS = macrolide, Fos = Fosfomycin, Phe = Phenicol

[1] MS7884 – IMP+ *E. cloacae* isolated from RBWH 2015

#### Phenotypic observations may relate to single nucleotide polymorphisms (SNPs)

Two isolates from Patient 3 exhibited *in vitro* phenotypic differences in adherence, suggesting that the respective SNPs may relate to within-host adaptation. Both isolates (MS7892 and MS7893) exhibited reduced pellet formation after centrifugation and resuspended more readily in solution compared to other cultures. This phenotype may have been advantageous for systemic infection, as both of these isolates were retrieved from wound and blood sites. Investigation into genotypic difference attributable to this phenotype found a non-synonymous SNP in *rcsB* (MS7884_3276) (Supplementary dataset S1), a well-known gene encoding a transcriptional regulatory protein that has previously been associated with colanic acid and biofilm production, and has been shown to provide protection from the bactericidal effect of serum during bloodstream infection [12, 13].

#### Patient 3 isolates are missing ~25 kb phage region

To improve our understanding of the transmission dynamics of this outbreak, we sought to identify differences in the pan-genome of these strains. Whole genome comparisons determined all 10 2015 RBWH isolates to have essentially identical genome content when compared to Ecl1. The exception was a ~25 kb region missing in all strains isolated from patient 3 (MS7890-MS7893), found to contain predicted phage-related proteins (Figure S1). Comparison of this region to the PHAST prophage database [8] found *Escherichia* phage HK639 (NC_016158.1) to be the closest match (35% coverage, 88% identity). This ~25 kb region represents only half of the full length of phage HK639 (~50 kb), and is present in Ecl1 and patient 1 and 2 isolates. It is unclear if the loss of this phage region could increase the fitness or virulence of the *E. hormaechei* variant infecting patient 3. Although the prophage region is likely to have been lost during the infection of patient 3, we cannot rule out that carbapenemase-producing *E.*

*hormaechei* with and without the phage co-exist within a common environmental source in the ICU.

#### AmpC is unlikely to contribute to reduced carbapenem susceptibility in the 2015 RBWH isolates

Chromosomally encoded *ampC* was detected in all 10 2015 RBWH *E. hormaechei* isolates (Table 1). While *ampC* is usually repressed and present at low levels in cells, SNPs resulting in derepression of *ampC* have been shown to increase resistance to beta-lactams [14, 15]. As such, we compared the nucleotide and amino acid sequence of our strains against several previously described SNPs in the regulatory genes *ampD*, *ampR* and *ampG* to predict *ampC* overexpression within our isolates.

All isolates contained 100% identical AmpC genes at both nucleotide and amino acid levels. The conserved AmpC residues Ser-64, Lys-67, Tyr-150, Asn-152, Lys-315 and Ala-318 [14] were all found to be present. Nucleotide and amino acid comparisons of AmpD between all 10 isolates were found to be 100% identical, with none of the previously described SNPs [16-18]. Amino acid comparison of AmpR to *E. cloacae* MNH1 [19] was 92% identical, with the highly conserved residues Arg-86, Gly-102, Ser-35, Tyr-264 and Asp-135 present in our strains. Comparison of AmpG to previous literature again found no obvious evidence to suggest heightened AmpC production [20-22]. We also checked the region upstream of *ampC* for the presence of insertion sequences (IS), as IS-driven overexpression of *ampC* has been previously reported [23-25]. Again, we found no evidence of IS upstream of *ampC*. We also found no evidence of plasmid-encoded *ampC*.

Additionally, observation of isolates grown with and without 2 ug/mL meropenem selection in LB found that colonies that lost the $bla_{\text{IMP-4}}$ carbapenemase were unable to grow under the meropenem selective pressure, further suggesting that the *bla*$_{IMP-4}$ carbapenemase is the major driver of carbapenem resistance in these strains, and that *ampC* expression has little or no effect.

#### Loss of ~34 kb region in MS7889 is mediated by homologous recombination

Compared to all other *E. hormaechei* from the 2015 RBWH isolates, MS7889 (patient 2, isolate 4) was found to have lost a number of antibiotic resistance genes, including *bla*$_{IMP-4}$. SMRT sequencing of the *E. hormaechei* isolate MS7884 (patient 1, isolate 1) enabled full resolution of a 330,060 bp IncHI2 plasmid harbouring a ~55 kb MDR region that encompassed the majority of antibiotic resistance genes. Analysis of this region and comparison to the draft assembly for MS7889 determined that the most likely mechanism of loss of *bla*$_{IMP-4}$ and other antibiotic resistance genes is homologous recombination between two near identical genes (*aacA4* and *aac(6')-Ib-cr),* resulting in the loss of a ~34 kb region. Nucleotide comparison of the two genes from MS7884 and the single *aac(6')-Ib-cr* gene from MS7889 identified only 3 SNPs between these genes (Figure S2).

Due to the similarity between *aacA4* and *aac(6')-Ib-cr*, these genes appear as a single contig in the Illumina draft assemblies. As such, these genes have only been characterised in the complete genome for MS7884, and have been left as *aac(6')-Ib-cr* in Table 1 for the remaining Illumina draft assemblies.

#### Unique SNPs in *aac(6')-Ib-cr* could result in increased resistance in MS7889

The carbapenem-sensitive isolate MS7889 was found by Vitek2 to have increased resistance to the aminoglycosides tobramycin and amikacin, despite the loss of several other antibiotic resistance genes. Comparison of the aminoglycoside resistance gene *aac(6')-Ib-cr* between the SMRT sequenced MS7884 reference and the MS7889 draft assembly identified 3 non-synonymous SNPs (Figure S3). A 329T SNP encoding Leucine was detected using MARA (7), which has previously been associated with increased amikacin resistance (28). The remaining two SNPs could further contribute to the observed increase in resistance to tobramycin and amikacin.

#### Whole genome sequencing identifies two plasmids carried in RBWH isolates

Illumina whole genome sequencing of the 10 RBWH *E. hormaechei* strains directly from clinical agar plates identified two large plasmids. Initial sequencing of isolate MS7884 (grown from a single colony) using PacBio SMRT sequencing was only able to identify one plasmid; a 126,208 bp un-typeable plasmid (pMS7884B) that lacked all of the previously identified antibiotic resistance genes determined from the Illumina sequencing data, including the $bla_{IMP-4}$ carbapenemase. Repeat PacBio SMRT sequencing of MS7884 (grown from a single colony with 2 µg/ml meropenem selection to avoid loss of $bla_{IMP-4}$) enabled the successful resolution of a large, 330,060 bp IncHI2 plasmid (pMS7884A) that was found to harbour all previously identified antibiotic resistance genes. However, pMS7884B was not present in this subsequent sequencing run. We therefore sought to determine whether these two plasmids were incompatible, and whether our clinical samples were comprised of a mixed population of carbapenemase-producing and non-producing *E. hormaechei.*

Glycerol stock of the original MS7884 clinical sample was streaked onto Mueller-Hinton (MH) agar and grown overnight at 37°C. 100 colonies were pick and patched onto both plain MH agar and MH agar with 1 µg/mL meropenem and grown overnight at 37°C. 15 of 100 colonies did not grow on the meropenem plate, suggesting loss of pMS7884A or $bla_{IMP-4}$. Colony PCR determined 8 of the 15 colonies retained pMS7884A, but had lost $bla_{IMP-4}$. The remaining 7 colonies appeared to have lost pMS7884A entirely.

Colony PCR of the 100 single colonies was performed using primers designed to target pMS7884B. Of 100 colonies, only 1 appeared to have lost the pMS7884B. This confirmed that these two plasmids are not incompatible and that loss of one of the two plasmids in MS7884 is not uncommon and independent plasmid loss likely occurred prior to PacBio SMRT sequencing.

#### pMS7884A and pMS7884B comparisons to published genomes

The complete ~55kb MS7884 MDR region shares the most similarity to a region identified in pEl1573, an IncL/M plasmid carrying *bla*$_{IMP-4}$ (isolated in Sydney 2012), with 99% nucleotide identity across 71% of the ~55 kb MDR region (Figure 4B) [26]. pMS7884A shares 86% of its IncHI2 backbone (with 99% nucleotide identity) to the previously reported IMP-producing pEC-IMPQ (GenBank: EU855788.1) plasmid isolated from China before 2009 [27]. The most closely related plasmid to pMS7884B in the NCBI database was the IncFII plasmid pECL_A (GenBank: CP001919.1) from *E. cloacae* subsp. cloacae ATCC 13047 [28], with 64% query coverage at 99% identity (Figure S4).

#### Summary of methylomes in complete PacBio genomes

Sequencing two single plasmid derivatives of the same isolate with PacBio SMRT sequencing enabled us to analyse the contribution of each plasmid to the genome-wide methylation status of every nucleotide (i.e. the methylome). Table S5 summarises the methylated motifs in MS7884A (with the ~330 kb IncHI2 plasmid) and MS7884B (with the ~126 kb untypeable plasmid), as determined using the SMRT Analysis Suite. Only one difference was determined between the two strains: an additional m6A methylated motif in MS7884B (ACCTRGCA).

**Table S5: Summary of recognition motifs[1] in MS7884A and MS7884B**

| Motif string | Modification type | Methylated (%) | Number Detected | Number in Genome | Mean Score (Qmod) | Mean IPD Ratio |
|---|---|---|---|---|---|---|
| **MS7884A** | | | | | | |
| GATC | m6A | 98.2 | 50733 | 51688 | 123.40524 | 4.0679536 |
| CCYAN$_9$TGAY | m6A | 98.0 | 450 | 459 | 122.58444 | 4.87911 |
| RTCAN$_9$TRGG | m6A | 97.4 | 447 | 459 | 121.434006 | 4.6271815 |
| CAGCNAC | m6A | 97.0 | 4521 | 4662 | 112.2818 | 3.8938332 |
| **MS7884B** | | | | | | |
| GATC | m6A | 99.1 | 49412 | 49856 | 238.59088 | 4.6236324 |
| RTCAN$_9$TRGG | m6A | 99.3 | 446 | 449 | 235.08296 | 5.354307 |
| CCYAN$_9$TGAY | m6A | 99.3 | 446 | 449 | 237.71748 | 5.4802275 |
| CAGCNAC | m6A | 98.1 | 4445 | 4530 | 218.76625 | 4.31024 |
| ACCTRGCA | m6A | 55.9 | 133 | 238 | 97.1203 | 1.9882706 |

[1] Probable false positives (modified bases and motifs with a Qmod < 80) are not shown

Searching this motif against REBASE PacBio motifs found 9 matches to previously identified PacBio motifs from *Salmonella* and *Enterobacter* species (CP015024, CP017087, CP017186, CP017187, CP017180, CP017179, CP017181, CP016012, CP016357, CP015923). As of yet an enzyme has not been determined for this motif.

To identify the causative methyltransferase (MTase) for this motif, sequences for the MS7884 chromosome, pMS7884A plasmid and pMS7884B plasmid were compared against the REBASE Gold Standard Database using BLASTn to determine MTase genes carried in the genome (Table S6). No previously identified MTase was found in pMS7884B when compared to the Rebase Gold Standard database suggesting the presence of a novel MTase on pMS7884B. Only one putative MTase gene was annotated on the plasmid: MS7884_pB0001 – a putative DNA adenine MTase (position 109-801 bp). Searches of the Conserved Domain Database at NCBI revealed that the encoded protein matched the Pfam DNA methylase domain (pfam01555; residues 21-203, E-value = 1.21e-41) and a more specific CDD domain with a putative methylase function (PRK13699; 1-219, E-value = 4.36e-44). As this is the only annotated MTase encoded on pMS7884B it is possible that MS7884_pB0001 encodes a novel MTase that accounts for the additional methylation detected at ACCTRGCA. However, we note that methylation was relatively low frequency (55.9% of sites), the mean score and IPD ratios are also low and the motif is not typical of adenine DNA MTases of any type, supporting the contention that MS7884_pB0001 may encode a m5C methylase that actually methylates CCWGG. Further work is required to elucidate the MTase properties (if any) of MS7884_pB0001.

**Table S6: Summary of methyltransferases**

| Strain | Hit[1] | Percentage identity (nt) | Query coverage | Locus Tag | Location |
|---|---|---|---|---|---|
| **MS7884A (chromosome)[2]** | | | | | |
| | M.EclNIH2 Dam GATC 813nt | 97% | 789/813 | MS7884_2116 | 2249369..2250181 |
| | M.Csa8155I GAANNNNNNntAAA 1563nt | 90% | 672/744 | MS7884_1344 | 1405871..1406629 |
| **pMS7884A (IncHI2)** | | | | | |
| | M.Sen 2050ORF235P GATC 813nt | 99% | 806/813 | MS7884_pA0268 | 227341..228153 |
| **pMS7884B (untypeable)** | | | | | |
| | MS7884_pB0001[3] | - | - | MS7884_pB0001 | 109..801 |

[1] Hits > 50 bp coverage, ≥ 90% identity with blastn against REBASE gold standard database (redundant hits excluded) downloaded 30/12/16
[2] MS7884A and MS7884B have identical gene content on the chromosome
[3] Putative methyltransferase identified on pMS7884B untypeable plasmid

#### Continued surveillance using Nanopore MinION sequencing

Since 2015 there has been continued surveillance of *bla*$_{IMP-4}$ positive Enterobacteriaceae in RBWH. In 2017, a *bla*$_{IMP-4}$ positive *E. hormaechei* was isolated from a patient within the Hematology ward (MS14449). We used Nanopore MinION sequencing to rapidly determine whether this isolate was related to the previous ST90 *E. cloacae* outbreak. Using the MinION sequencing data alone we were able to create draft assemblies for the chromosome (~5 Mbps) and two plasmids (~382 kb and ~145 kb). However, due to a high abundance of errors we were unable to accurately determine the ST of MS14449. The larger plasmid in MS14449 appeared identical to pMS7884A based on comparison using ACT (5). The exception was a large region (position 22250-143210 in pMS7884A) that appeared to be inverted (Figure S7). This region was flanked by IS*26* insertion sequences, which likely facilitated this inversion. The inverted region also encompasses the tetracycline and chloramphenicol resistance genes from the MDR region described in pMS7884A. Comparison of the smaller plasmid in MS14449 to pMS7884B using BRIG identified a shared core region, but large differences throughout the rest of the plasmid (Figure S8).

In order to determine if MS14449 was clonally related to the 2015 outbreak *E. hormaechei*, we built a tree using Parsnp (v1.2) to contextualise the draft assembly for MS14449 against all other publicly available complete *E. cloacae* complex strains from NCBI (accessed 2/11/2017). This analysis placed MS14449 outside of the ST90 outbreak *E. hormaechei* cluster, indicating that it was unlikely to be related at the strain level (Figure S9). MS14449 was later confirmed as a different ST (ST175) using Illumina sequencing.

A *Klebsiella pneumoniae* isolate was also recovered from the same patient as MS14449 and was found to be $bla_{\mathrm{IMP-4}}$ positive (MS14448). Illumina sequencing of this isolate and nucleotide comparison to pMS7884A indicated that it likely carries a very similar plasmid, with the exception of a small section of the MDR, which it appears to have lost (Figure S10).

#### Environmental surveillance in the hospital

**Table S7: Isolates cultured from environmental surveillance swabs**

| Isolate | Associated environmental sample | Cultured species ID | ST | IncHI2 plasmid? |
|---|---|---|---|---|
| M87132 | R5505 | *Klebsiella oxytoca*[1] IMP4+ | 88 | N |
| M87133 | R5506 | *Enterobacter cloacae* complex IMP4+ | 254 | Y |
| M87134 | R5514 | *Enterobacter cloacae* complex | 830 | Y |
| M87135 | R5521 | *Leclercia adecarboxylata* IMP4+ | n/a | Y |

[1] Whole genome sequencing re-identified this isolate as *Klebsiella michiganensis*

**Table S8: MLST alleles detected in MAGs from samples with *E. hormaechei* detected based on MASH**

| MAG*/Isolate | dnaA | fusA | gyrB | leuS | pyrG | rplB | rpoB |
|---|---|---|---|---|---|---|---|
| ST90 *E. hormaechei* (MS7884) | 58 | 37 | 4 | 6 | 42 | 4 | 25 |
| R5514 | ~58 | ~37 | 62? | 6 | 42,67 | 48? | 25 |
| R5506 | - | - | - | - | - | - | 6? |
| R5522 | 129? | 8? | 21? | 41? | 15? | 79? | - |
| R5537 | 58 | ~37 | 62? | ~6 | 42,67 | 4 | 25 |

#### Supplementary Figures

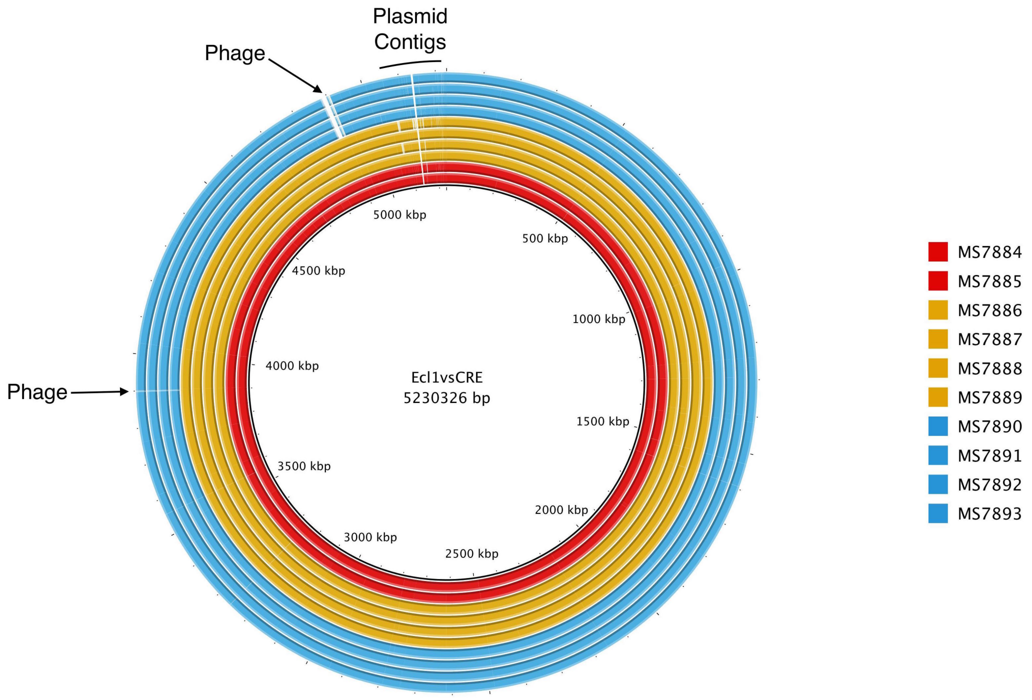


**Figure S1: Whole genome comparison of RBWH draft assemblies using BRIG:** Draft assemblies of all RBWH isolates were compared against the Ecl1 draft reassembly (*E. cloacae*, 2013 RBWH) using the blast comparison tool BRIG [6]. Whole genome comparison identified two main differences between the patient 3 isolates (blue) and the patient 1 and 2 isolates (red and gold, respectively). All patient 3 isolates were missing a ~25 kb region identified as containing phage-related genes.

```
                        1
MS7884_aacA4            ---------- ---------- ---------- ---------- -----GTGAC CAACAGCAAC
MS7889_aac(6')-Ib-cr    ATGAGCAACG CAAAAACAAA GTTAGGCATC ACAAAGTACA GCATCGTGAC CAACAGCAAC
MS7884_aac(6')-Ib-cr    ATGAGCAACG CAAAAACAAA GTTAGGCATC ACAAAGTACA GCATCGTGAC CAACAGCAAC

                       61
MS7884_aacA4            GATTCCGTCA CACTGCGCCT CATGACTGAG CATGACCTTG CGATGCTCTA TGAGTGGCTA
MS7889_aac(6')-Ib-cr    GATTCCGTCA CACTGCGCCT CATGACTGAG CATGACCTTG CGATGCTCTA TGAGTGGCTA
MS7884_aac(6')-Ib-cr    GATTCCGTCA CACTGCGCCT CATGACTGAG CATGACCTTG CGATGCTCTA TGAGTGGCTA

                      121
MS7884_aacA4            AATCGATCTC ATATCGTCGA GTGGTGGGGC GGAGAAGAAG CACGCCCGAC ACTTGCTGAC
MS7889_aac(6')-Ib-cr    AATCGATCTC ATATCGTCGA GTGGTGGGGC GGAGAAGAAG CACGCCCGAC ACTTGCTGAC
MS7884_aac(6')-Ib-cr    AATCGATCTC ATATCGTCGA GTGGTGGGGC GGAGAAGAAG CACGCCCGAC ACTTGCTGAC

                      181
MS7884_aacA4            GTACAGGAAC AGTACTTGCC AAGCGTTTTA GCGCAAGAGT CCGTCACTCC ATACATTGCA
MS7889_aac(6')-Ib-cr    GTACAGGAAC AGTACTTGCC AAGCGTTTTA GCGCAAGAGT CCGTCACTCC ATACATTGCA
MS7884_aac(6')-Ib-cr    GTACAGGAAC AGTACTTGCC AAGCGTTTTA GCGCAAGAGT CCGTCACTCC ATACATTGCA

                      241
MS7884_aacA4            ATGCTGAATG GAGAGCCGAT TGGGTATGCC CAGTCGTACG TTGCTCTTGG AAGCGGGGAC
MS7889_aac(6')-Ib-cr    ATGCTGAATG GAGAGCCGAT TGGGTATGCC CAGTCGTACG TTGCTCTTGG AAGCGGGGAC
MS7884_aac(6')-Ib-cr    ATGCTGAATG GAGAGCCGAT TGGGTATGCC CAGTCGTACG TTGCTCTTGG AAGCGGGGAC

                      301
MS7884_aacA4            GGATGGTGGG AAGAAGAAAC CGATCCAGGA GTACGCGGAA TAGACCAGTC ACTGGCGAAT
MS7889_aac(6')-Ib-cr    GGATGGTGGG AAGAAGAAAC CGATCCAGGA GTACGCGGAA TAGACCAGTT ACTGGCGAAT
MS7884_aac(6')-Ib-cr    GGAAGGTGGG AAGAAGAAAC CGATCCAGGA GTACGCGGAA TAGACCAGTT ACTGGCGAAT

                      361
MS7884_aacA4            GCATCACAAC TGGGCAAAGG CTTGGGAACC AAGCTGGTTC GAGCTCTGGT TGAGTTGCTG
MS7889_aac(6')-Ib-cr    GCATCACAAC TGGGCAAAGG CTTGGGAACC AAGCTGGTTC GAGCTCTGGT TGAGTTGCTG
MS7884_aac(6')-Ib-cr    GCATCACAAC TGGGCAAAGG CTTGGGAACC AAGCTGGTTC GAGCTCTGGT TGAGTTGCTG

                      421
MS7884_aacA4            TTCAATGATC CCGAGGTCAC CAAGATCCAA ACGGACCCGT CGCCGAGCAA CTTGCGAGCG
MS7889_aac(6')-Ib-cr    TTCAATGATC CCGAGGTCAC CAAGATCCAA ACGGACCCGT CGCCGAGCAA CTTGCGAGCG
MS7884_aac(6')-Ib-cr    TTCAATGATC CCGAGGTCAC CAAGATCCAA ACGGACCCGT CGCCGAGCAA CTTGCGAGCG

                      481
MS7884_aacA4            ATCCGATGCT ACGAGAAAGC GGGGTTTGAG AGGCAAGGTA CCGTAACCAC CCCAGATGGT
MS7889_aac(6')-Ib-cr    ATCCGATGCT ACGAGAAAGC GGGGTTTGAG AGGCAAGGTA CCGTAACCAC CCCAGATGGT
MS7884_aac(6')-Ib-cr    ATCCGATGCT ACGAGAAAGC GGGGTTTGAG AGGCAAGGTA CCGTAACCAC CCCATATGGT

                      541
MS7884_aacA4            CCAGCCGTGT ACATGGTTCA AACACGCCAG GCATTCGAGC GAACACGCAG TGATGCCTAA
MS7889_aac(6')-Ib-cr    CCAGCCGTGT ACATGGTTCA AACACGCCAG GCATTCGAGC GAACACGCAG TGATGCCTAA
MS7884_aac(6')-Ib-cr    CCAGCCGTGT ACATGGTTCA AACACGCCAG GCATTCGAGC GAACACGCAG TGATGCCTAA
```

**Figure S2: Nucleotide alignment between MS7884 *aac(6')-Ib-cr*, MS7884 *aacA4* and MS7889 *aac(6')-Ib-cr* gene**: red boxes indicate SNP positions

```
                      1
MS7884_aacA4          ---------- -----VTNSN DSVTLRLMTE HDLAMLYEWL NRSHIVEWWG GEEARPTLAD
MS7889_aac(6')-1b-cr  MSNAKTKLGI TKYSIVTNSN DSVTLRLMTE HDLAMLYEWL NRSHIVEWWG GEEARPTLAD
MS7884_aac(6')-1b-cr  MSNAKTKLGI TKYSIVTNSN DSVTLRLMTE HDLAMLYEWL NRSHIVEWWG GEEARPTLAD

                     61                                      *                *
MS7884_aacA4          VQEQYLPSVL AQESVTPYIA MLNGEPIGYA QSYVALGSGD GWWEEETDPG VRGIDQSLAN
MS7889_aac(6')-1b-cr  VQEQYLPSVL AQESVTPYIA MLNGEPIGYA QSYVALGSGD GWWEEETDPG VRGIDQLLAN
MS7884_aac(6')-1b-cr  VQEQYLPSVL AQESVTPYIA MLNGEPIGYA QSYVALGSGD GRWEEETDPG VRGIDQLLAN

                    121                                                           *
MS7884_aacA4          ASQLGKGLGT KLVRALVELL FNDPEVTKIQ TDPSPSNLRA IRCYEKAGFE RQGTVTTPDG
MS7889_aac(6')-1b-cr  ASQLGKGLGT KLVRALVELL FNDPEVTKIQ TDPSPSNLRA IRCYEKAGFE RQGTVTTPDG
MS7884_aac(6')-1b-cr  ASQLGKGLGT KLVRALVELL FNDPEVTKIQ TDPSPSNLRA IRCYEKAGFE RQGTVTTPYG

                    181
MS7884_aacA4          PAVYMVQTRQ AFERTRSDA
MS7889_aac(6')-1b-cr  PAVYMVQTRQ AFERTRSDA
MS7884_aac(6')-1b-cr  PAVYMVQTRQ AFERTRSDA
```

**Figure S3: Multiple sequence alignment of *aacA4* and *aac(6')-Ib-cr* amino acid sequence in MS7884 and MS7889:** red asterisks indicate amino acid change sites

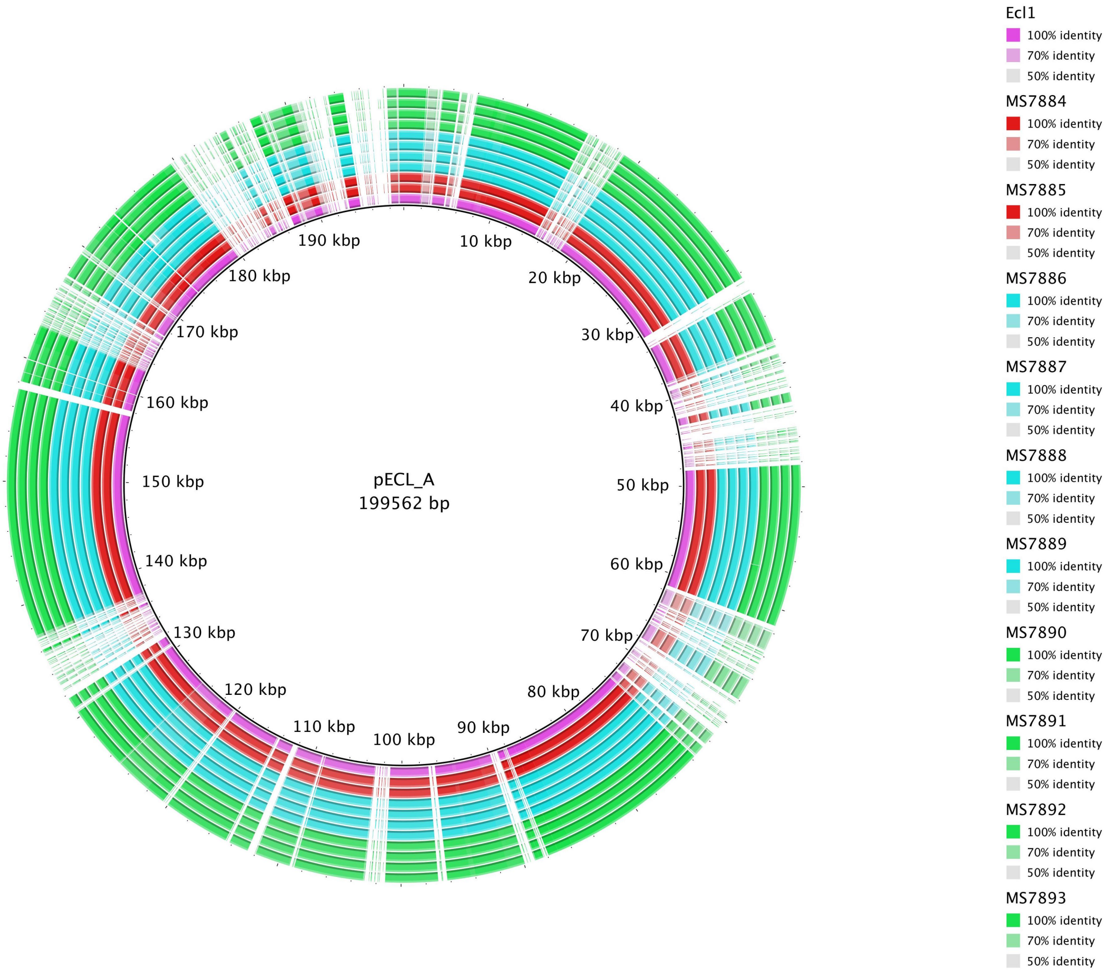


**Figure S4: BRIG comparison of 2015 RBWH draft assemblies to pECL_A (GenBank: CP001919.1) from E. cloacae subsp. cloacae ATCC 13047:** pMS7884B was found to be most closely related to pECL_A from *E. cloacae* subsp. cloacae ATCC 13047 (BLASTn; 64% query coverage at 99% identity). Draft assemblies for the 2013 strain Ecl1 (pink), 2015 Patient 1 strains (red), 2015 Patient 2 strains (light blue), 2015 Patient 3 strains (green) were compared against pECL_A using BRIG [6].

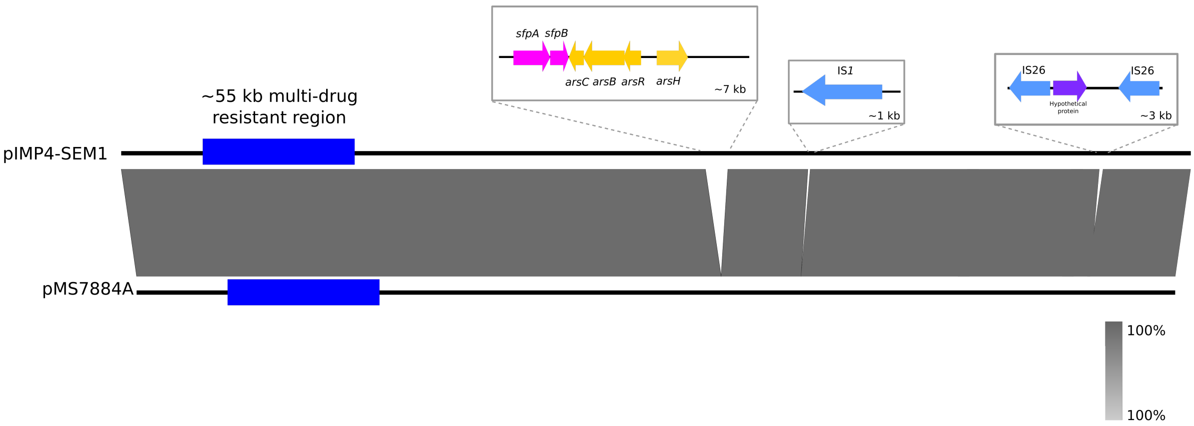


**Figure S5: Easyfig comparison of pIMP4-SEM1 and pMS7884A:** Black lines represent the complete plasmids pIMP4-SEM1 and pMS7884A. Grey blocks indicate nucleotide identity between the two plasmids. Both plasmids were found to be near identical, carrying the same large ~55 kb multi-drug resistant region. Three additional regions were found in pIMP4-SEM1: a ~7 kb region carrying *sfpAB* and *arsCBRH*, an IS*1* inserted upstream of *trhR,* and an IS26-bound transposon carrying a hypothetical protein.

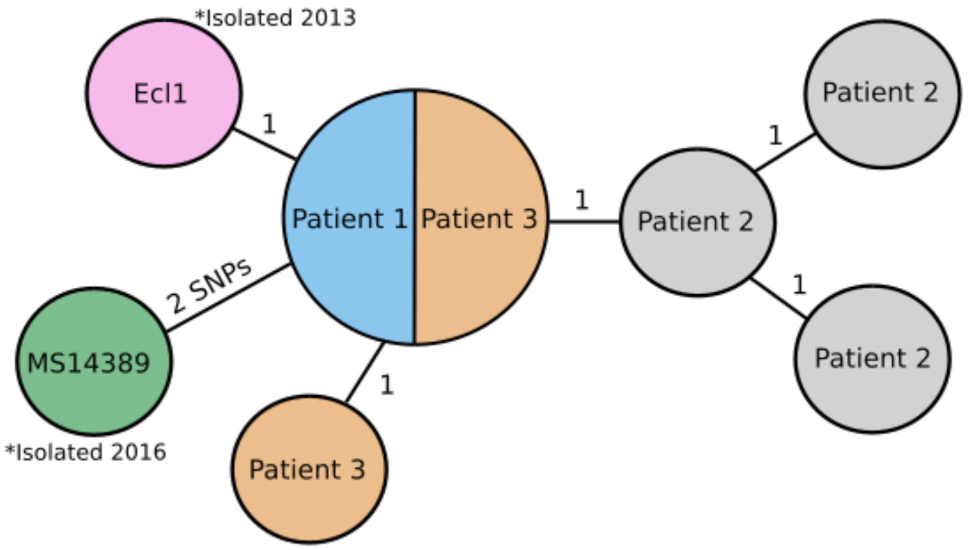


**Figure S6: Relationship matrix of outbreak ST90 *E. hormaechei* with MS14389**

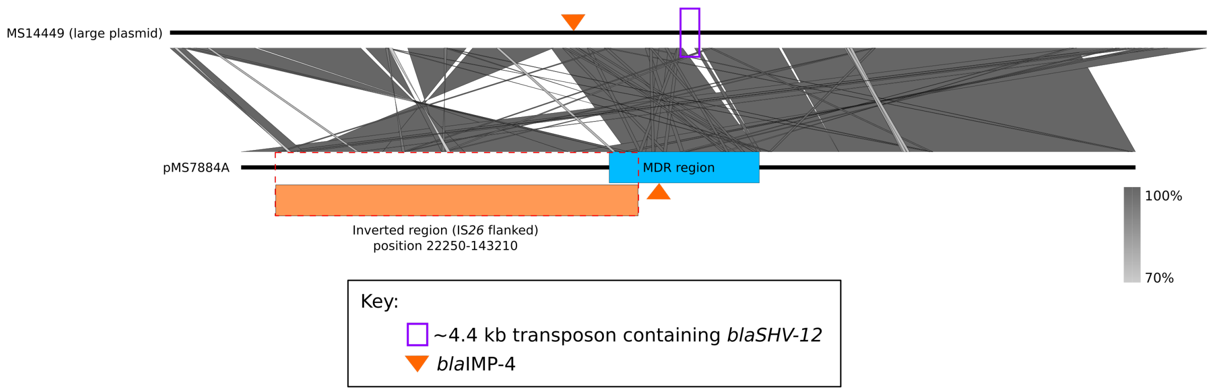


**Figure S7: Comparison of large plasmid from MS14449 and pMS7884A**

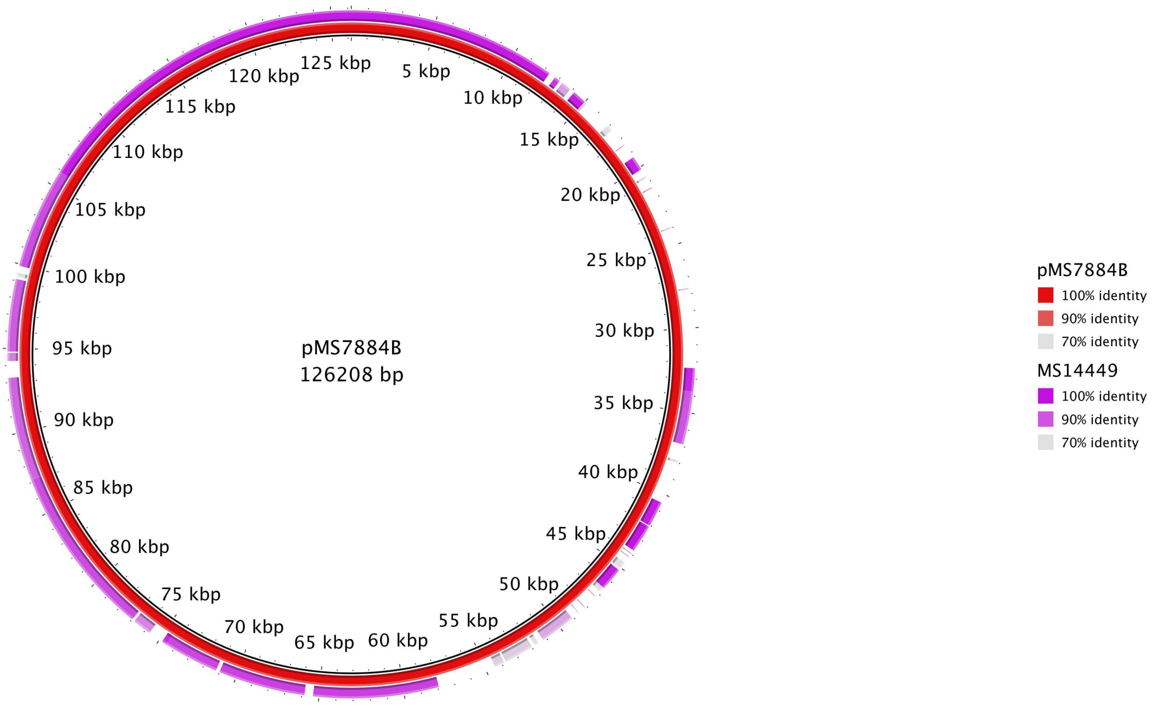


**Figure S8: BRIG comparison of pMS7884B and small plasmid from MS14449**

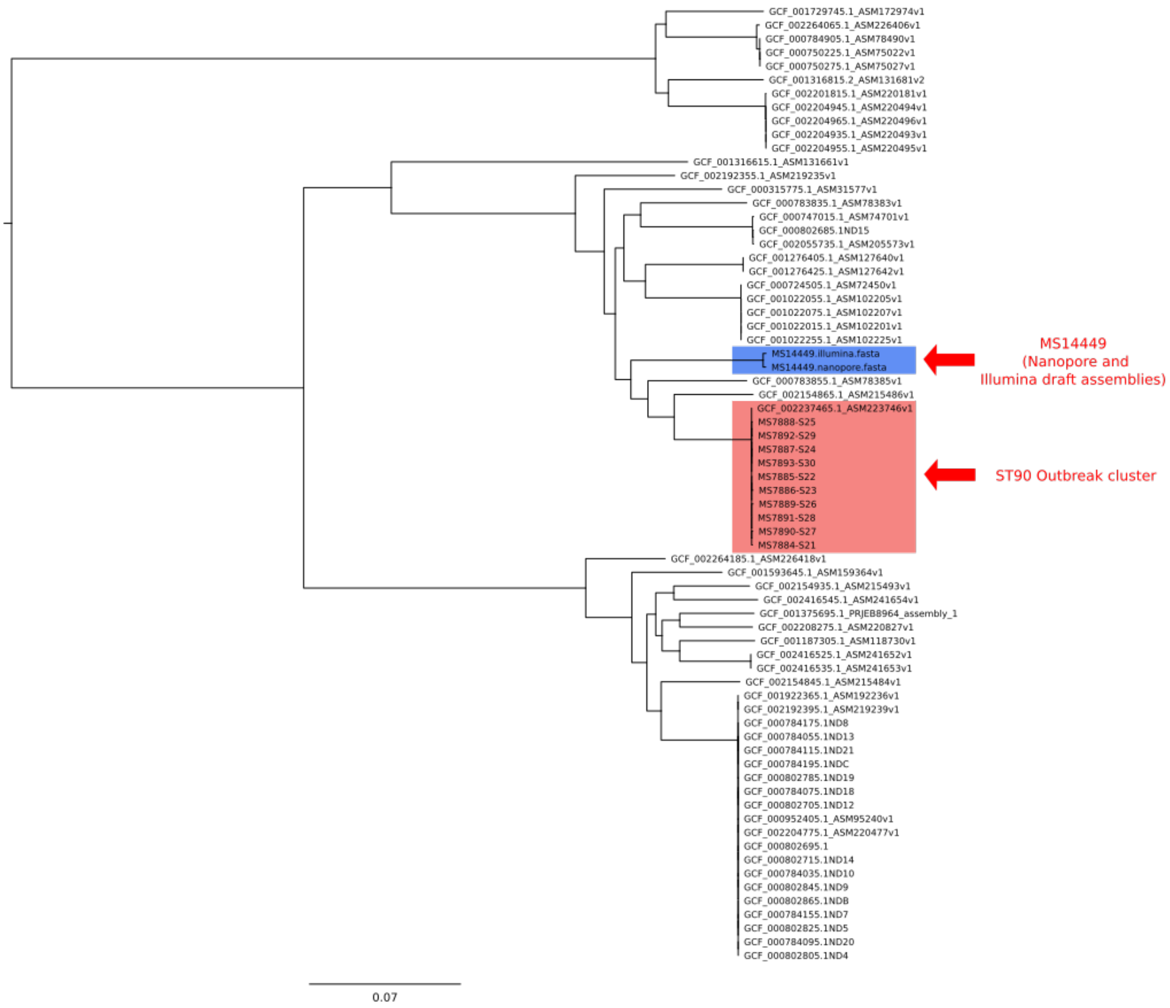


**Figure S9: Parsnp tree contextualising MS14449 Illumina and Nanopore draft assemblies against ST90 outbreak cluster and other complete publicly available *E. cloacae* complex strains**

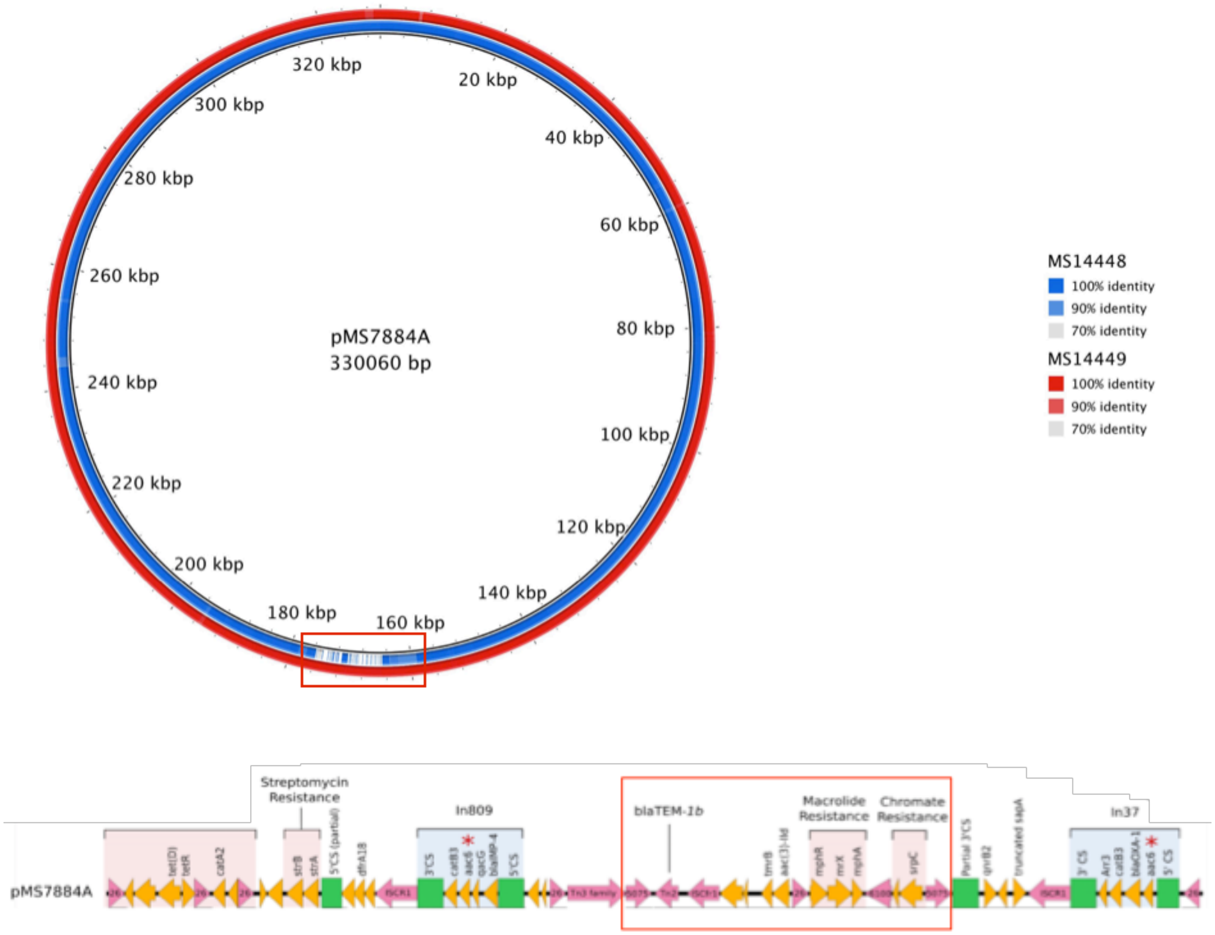


**Figure S10: BRIG comparison of MS14448 and MS14449 draft assemblies against pMS7884A:** red box indicates region missing from MS14448 *K. pneumoniae* IncHI2 plasmid

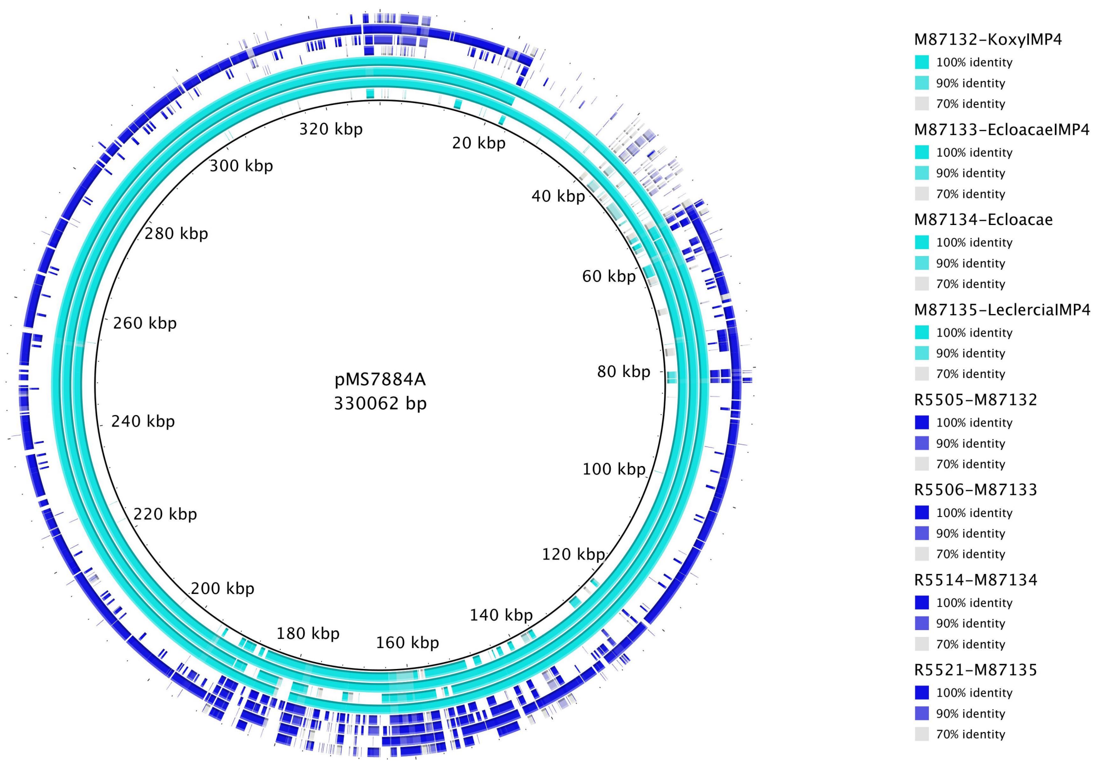


**Figure S11: BRIG comparison of isolate draft assemblies (light blue, see Table S7) and associated MAGs from metagenomic sequencing of associated environmental samples (dark blue) against IncHI2 plasmid pMS7884A.**

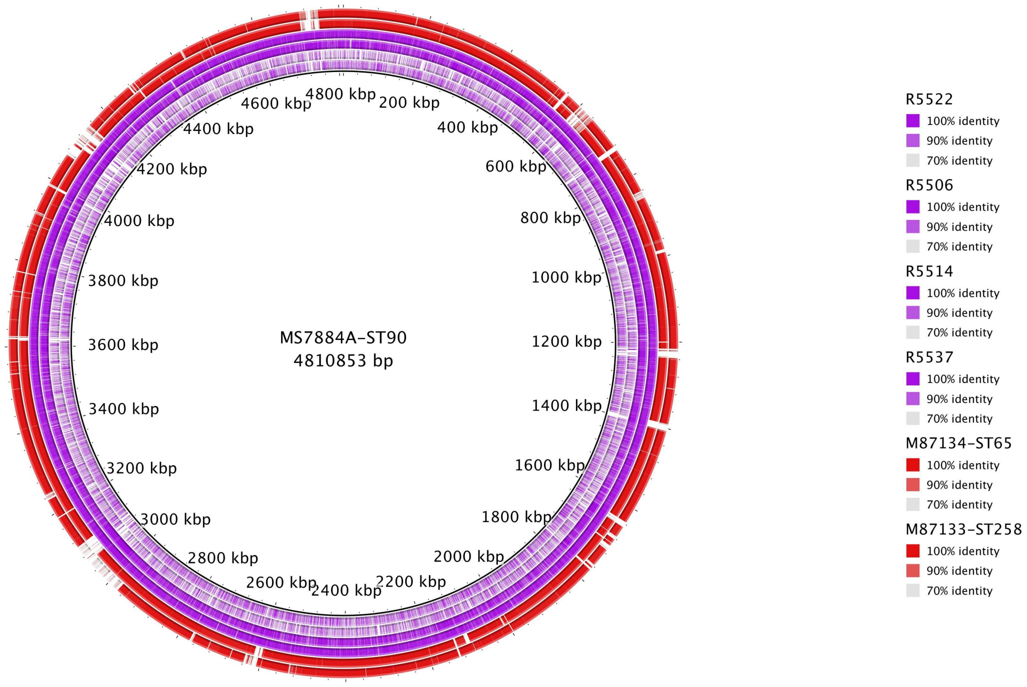


**Figure S12: BRIG of MAGs from samples with positive *E. hormaechei* (purple; as determined by MASH) against the chromosome of the ST90 *E. hormaechei* strain MS7884A. Red rings represent *E. cloacae* complex strains of different ST for comparison.** R5514 and R5537 appear to have high similarity to MS7884A.

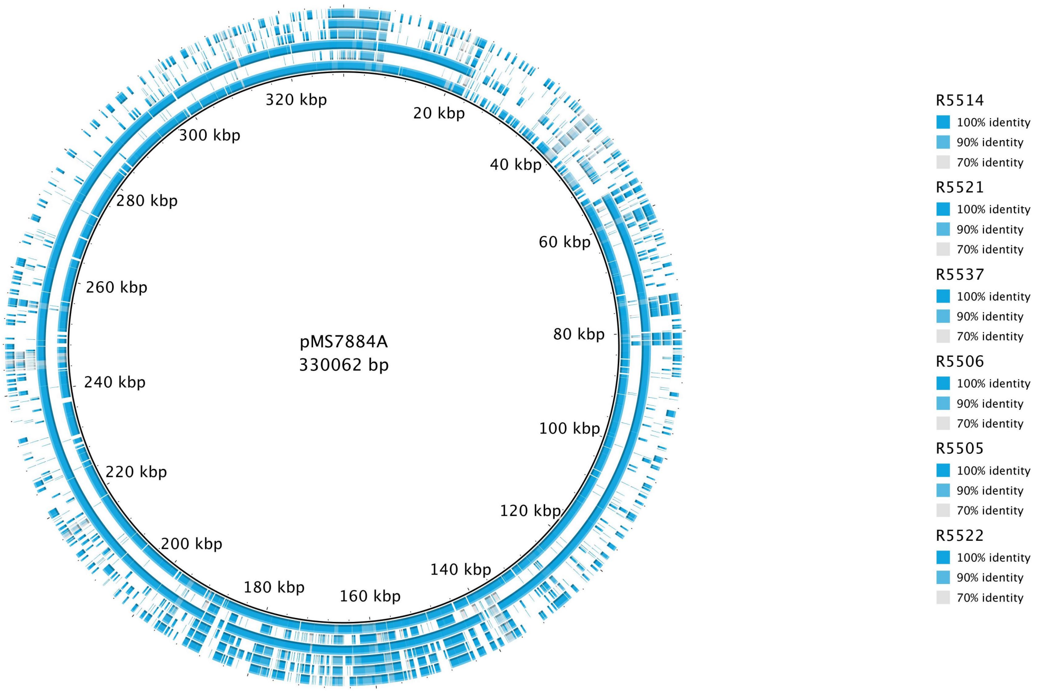


**Figure S13: BRIG of MAGs from samples with positive pMS7884A and/or the associated MDR region (as determined by MASH) against the IncHI2 plasmid pMS7884A.** Two samples (R5514 and R5537) appear to have high similarity to pMS7884A.
